# Integrated Profiling of Synaptic E/I Balance Reveals Altered Synaptic Organization and Phenotypic Variability in a Prenatal ASD Model

**DOI:** 10.1101/2025.06.08.658540

**Authors:** Mirei Matsumuro, Sotaro Ichinose, Hirohide Iwasaki

## Abstract

Proper regulation of excitation and inhibition (E/I balance) is essential for maintaining stable neural circuit function and flexible behavior. Disruptions of E/I balance have been implicated in a variety of neurodevelopmental and psychiatric disorders, including autism spectrum disorder (ASD). However, despite advances in connectomics and molecular neuroscience, how localized disruptions in E/I balance within cortical laminar architectures contribute to behavioral abnormalities remains to be elucidated. Here, we developed a standardized analysis pipeline that combines depth-aligned synaptic mapping, perspective and logarithmic transformations, and multivariate behavioral profiling. Applying this approach to a prenatal valproic acid (VPA) exposure model of ASD, we identified depth-specific disruptions of excitatory and inhibitory synaptic organization within the anterior cingulate cortex (ACC), alongside impairments in social behavior. Principal component analysis (PCA) integrating synaptic and behavioral parameters revealed convergent abnormalities that robustly distinguished VPA-exposed mice from wild-type (WT) controls. Furthermore, although still at an exploratory stage, our findings demonstrate that the robustness of this pipeline enables both the identification of resilient individuals and the quantification of intervention effects. Together, our findings establish a new cross-level analytical framework linking synaptic organization to organismal behavior and provide insights into circuit-level mechanisms underlying ASD-related phenotypes.

## Introduction

Proper regulation of excitation and inhibition (E/I balance) within neural circuits is fundamental for maintaining stable brain function and flexible behavioral control (Sohal & Rubenstein, 2019). At the cellular level, E/I balance governs neuronal excitability through spatiotemporal integration of synaptic inputs, ensuring appropriate action-potential generation (Isaacson & Scanziani, 2011; Spruston, 2008). At the network level, coordinated modulation of E/I balance sustains oscillatory activity that underlies higher cognitive and sensorimotor functions such as attention, working memory and emotional regulation (McKeon et al., 2024). Disruptions of E/I balance have been implicated in a wide range of neurodevelopmental and psychiatric disorders, including autism spectrum disorder (ASD), schizophrenia, and epilepsy (Gogolla et al., 2009; Liu et al., 2021; Righes Marafiga et al., 2021; Yizhar et al., 2011). Mechanistically, E/I balance is stabilized through homeostatic plasticity processes that dynamically adjust synaptic strength and intrinsic excitability to maintain circuit activity (Chen et al., 2022). Key molecular pathways include intracellular transport of AMPA receptors and adhesion molecules, GABA_A_ receptor clustering, and associated intracellular signaling cascades (Ichinose et al., 2015; Turrigiano, 2012; Welle & Smith, 2025). Despite these advances, how localized E/I imbalances within cortical laminar architectures are integrated and transformed into circuit-level and behavioral dysfunction remains to be elucidated.

ASD is a highly heterogeneous neurodevelopmental disorder, clinically characterized by persistent deficits in social communication and interaction, as well as restricted and repetitive patterns of behavior (American Psychiatric, 2013; Lord et al., 2020). Despite this phenotypic diversity, convergent evidence from genetic, molecular and imaging studies indicates that alterations in synaptic excitation–inhibition (E/I) balance represent a shared pathophysiological mechanism across many ASD aetiologies (Nelson & Valakh, 2015). Functional and structural imaging studies have revealed widespread connectivity alterations in ASD, with the anterior cingulate cortex (ACC) frequently reported as a hub involved in social cognition, emotional regulation and executive function (Dichter, 2012; Zerbi et al., 2021; Zhou et al., 2016).

Although advances in connectomics have clarified large-scale network abnormalities (Anastasiades & Carter, 2021; van den Heuvel & Sporns, 2019), we still know far less about how microcircuit-level disruptions propagate to affect global information processing and behavior, particularly at cellular and synaptic resolution (Lichtman & Denk, 2011; Luo, 2021; Shaw et al., 2021; Yang et al., 2016). Pinpointing the cortical layers and microdomains where E/I imbalance arises is crucial, because local structural alterations are likely to compromise long-range integration and the computational capabilities of brain networks (Allen & Morishita, 2024; Keijser & Sprekeler, 2022). Without resolving such microstructural pathology, our understanding of how neurodevelopmental disorders emerge from circuit dysfunction remains fundamentally incomplete.

Rodent models generated by prenatal exposure to valproic acid (VPA) provide a valuable platform for dissecting ASD-related circuit mechanisms (Nicolini & Fahnestock, 2018). VPA exposure during critical periods of neurodevelopment can produce offspring that exhibit core ASD-like behaviors, including social interaction deficits, increased repetitive behaviors, and, in some studies, sensory hypersensitivity and anxiety-like traits (Mabunga et al., 2015). Electrophysiological studies in these models have revealed alterations in both excitatory and inhibitory synaptic transmission, suggestive of circuit-level E/I imbalance (Qi et al., 2022). Nevertheless, direct histological mapping of synaptic organization across cortical layers—and its integration with behavioral profiling—has remained technically challenging. Given that cortical laminar architecture supports the segregation, integration and output of distinct information streams (Harris & Shepherd, 2015), resolving how synaptic E/I balance is disrupted across cortical depths is critical for understanding ASD pathophysiology.

Here, we developed a standardized pipeline that combines depth-aligned synaptic mapping, perspective and logarithmic transformations, and multivariate behavioral profiling. Applying this approach to the prenatal VPA model, we identified depth-specific disruptions in excitatory and inhibitory synaptic organization within the ACC and showed that these structural abnormalities converge with behavioral impairments in social interaction and working memory. Principal component analysis (PCA) integrating synaptic and behavioral parameters revealed convergent patterns that robustly distinguished VPA-exposed mice from wild-type (WT) controls. Together, our findings establish a cross-level analytical framework that links cortical circuit pathology to organism-level behavioral outcomes in neurodevelopmental disorders.

## Results

### VPA-exposed mice exhibit ASD-like behaviors including impaired social interaction and decreased anxiety

To systematically investigate how circuit-level abnormalities shape behavioral phenotypes in ASD models, we applied our newly developed analysis pipeline to mice prenatally exposed to VPA. This standardized pipeline combines depth-aligned synaptic mapping, perspective-logarithmic transformations, and integrated multivariate profiling of behavioral and synaptic parameters.

Male mice were weaned at postnatal week 3 and housed under conditions with or without voluntary exercise until 12 weeks of age, at which point behavioral testing, tissue fixation, immunohistochemistry, and laser microdissection (LMD)-based qPCR analyses were performed (Figure 1A). For subsequent molecular and histological analyses, ACC samples were collected from coronal sections centered around the anatomical landmark where the left and right corpus callosum converge. This anatomical reference enabled consistent layer-specific analyses across individuals, accommodating natural variation in brain morphology.

**Figure 1.**
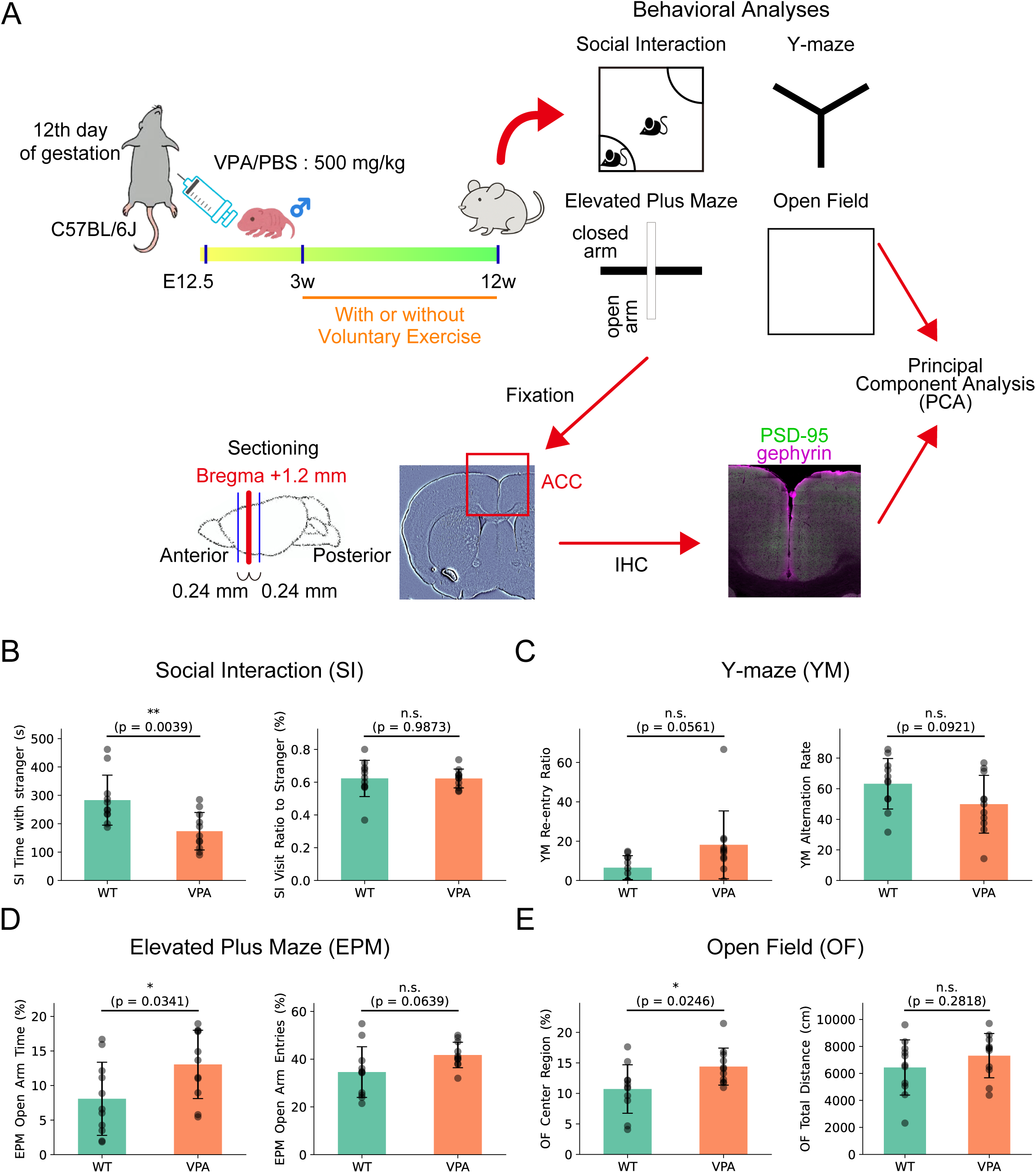
Behavioral analyses in WT and VPA-exposed mice. (A) Experimental timeline. Pregnant mice were administered valproic acid (500 mg/kg, intraperitoneally) on embryonic day 12.5 (E12.5). Mice were weaned at 3 weeks of age and subsequently housed with or without access to voluntary exercise. Behavioral analyses, histological analysis, and quantitative PCR (qPCR) were performed at 12 weeks of age. Immunohistochemical staining was performed on three coronal sections centered around Bregma +1.2 mm, spanning ±0.24 mm. Sections were selected based on the anatomical landmark of the fused corpus callosum rather than precise stereotaxic coordinates. Each quantitative analysis result was integrated using principal component analysis (PCA). (B) Summary of Social Interaction (SI) test. (C) Summary of Y-Maze (YM) test. (D) Summary of Elevated Plus Maze (EPM) test. (E) Summary of Open Field test. Data are presented as mean ± SD. WT (*n* = 11), VPA (*n* = 11). Statistical analysis was performed using Welch’s *t*-test. * *p* < 0.05; ** *p* < 0.01; n.s. not significant.

We first evaluated behavioral phenotypes in VPA-exposed mice using a series of established assays to assess social interaction, working memory, anxiety-related behaviors, and general locomotor activity (Figure 1A). VPA-exposed mice exhibited significant deficits in social interaction compared with WT controls, as indicated by reduced time in the social zone (WT: 283.1 ± 88.2 sec; VPA: 173.6 ± 66.0 sec; *p* = 0.0039), whereas the number of social approach events, assessed by the visit ratio to the stranger mouse, remained unchanged (WT: 0.623 ± 0.110; VPA: 0.622 ± 0.057; *p* = 0.9873), suggesting that social motivation was preserved (Figure 1B and Figure 1—figure supplement 1A). VPA-exposed mice displayed a trend toward reduced working-memory performance, with an increased re-entry ratio (WT: 6.5 ± 6.1 %; VPA: 18.1 ± 17.3 %; *p* = 0.0561) and a decreased alternation rate (WT: 63.3 ± 16.5 %; VPA: 49.9 ± 18.9 %; *p* = 0.0921), although neither reached statistical significance (Figure 1C). Anxiety-like behavior was decreased, as reflected by a higher proportion of time in the open arms of the elevated plus maze (EPM) (WT: 8.1 ± 5.3 %; VPA: 13.1 ± 4.9 %; *p* = 0.0341) (Figure 1D and Figure 1—figure supplement 1B) and a greater time in the center of the open field (WT: 10.7 ± 4.0 sec; VPA: 14.4 ± 3.0 sec; *p* = 0.0246) (Figure 1E and Figure 1—figure supplement 1C). Locomotor activity, measured as total distance traveled (WT: 6439.9 ± 2045.9 cm; VPA: 7315.6 ± 1640.4 cm; *p* = 0.282), did not differ between groups, indicating that the behavioral changes were not confounded by motor impairments. Although total travel distance in the Y-maze was significantly reduced, no major changes in locomotor activity were observed across other behavioral paradigms (Figure 1—figure supplement 1D–F). Therefore, the reduced distance in the Y-maze is likely a secondary reflection of impaired working memory rather than a locomotor deficit, suggesting that motor function remains largely intact in VPA-exposed mice.

These findings indicate that prenatal VPA exposure produces robust, selective alterations in social and anxiety-related behaviors while sparing general locomotion, providing a solid behavioral basis for probing the underlying synaptic and molecular mechanisms.

### Perspective transformation and logarithmic depth correction for accurate depth-resolved analysis in the ACC

Prenatal VPA exposure is a widely used model for studying ASD, but interpreting the resulting behavioral and molecular data is complicated by their variability. In this study, we focused on the ACC, whose irregular, curved anatomy presents challenges for depth-resolved synaptic quantification. To address this, we developed a standardized pipeline combining histogram matching (Figure 2—figure supplement 1A), perspective transformation, and logarithmic scaling (Figure 2A). The ACC was divided into small tiles for synapse counting, then computationally flattened to assign each tile a normalized depth position. These transformations reduced spatial and technical variability, enabling consistent depth-specific analyses across individuals.

**Figure 2.**
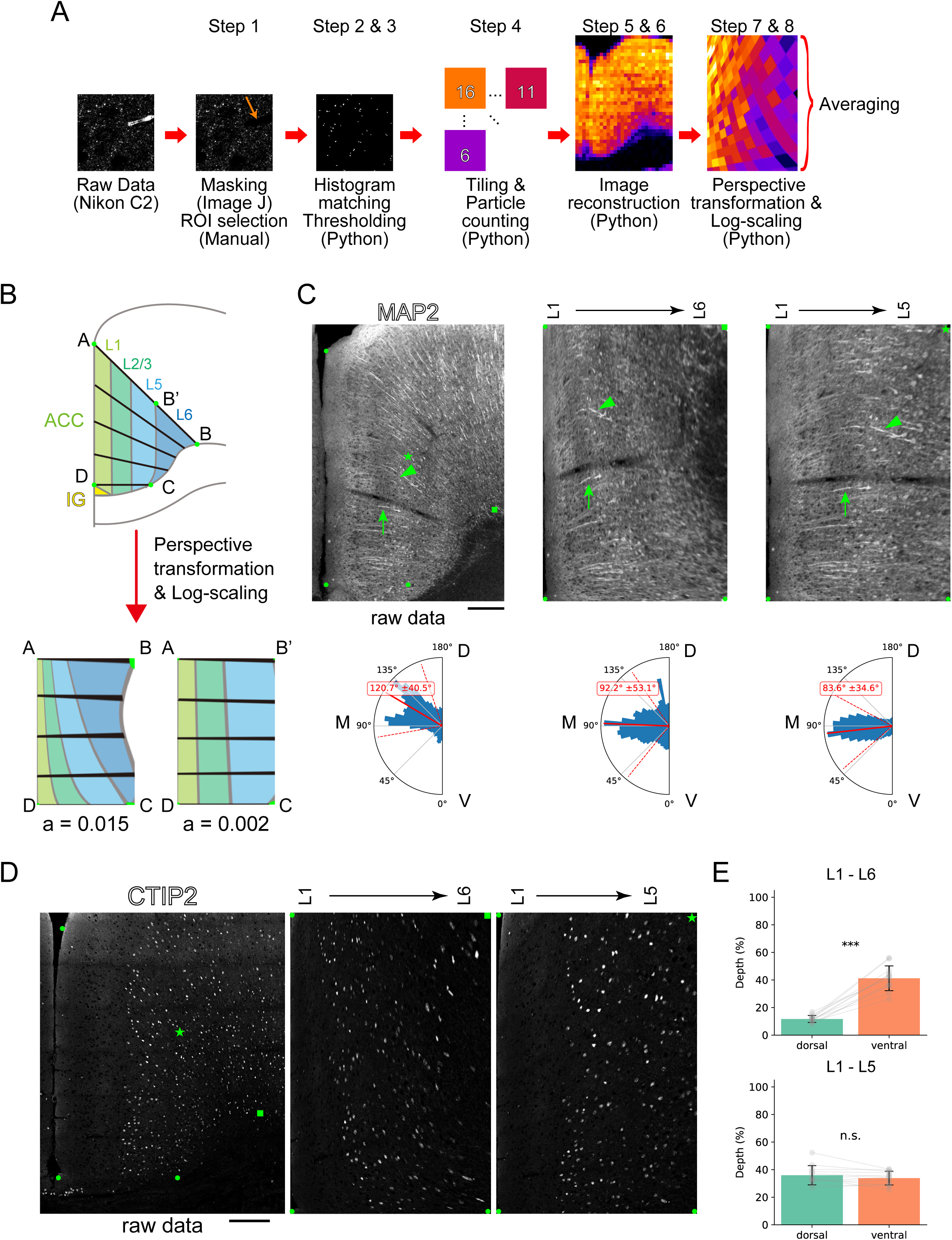
Perspective and log transformation for accurate depth-resolved analysis in the ACC. (A) Schematic of the standardized analysis pipeline. Confocal images were subjected to appropriate masking and ROI selection (Step 1). To reduce spatial and technical variability, histogram matching was applied, followed by Laplacian of Gaussian (LoG) filtering and thresholding (Steps 2 and 3). Images were then divided into small tiles for quantification (Step 4), and synapse maps were generated (Steps 5 and 6). Perspective transformation and logarithmic scaling were applied to computationally flatten the maps, and each tile was assigned a normalized depth value (Steps 7 and 8). (B) Model-based simulation of the transformation process. To detect layers 1–6, four points (A, B, C, D) were selected, and perspective transformation with log-scaling was applied using a = 0.015. To detect layers 1–5, four alternative points (A, B’, C, D) were selected with a = 0.002 for transformation and scaling. (C) MAP2 staining and the histogram showing the distribution of dendritic orientation and directionality from each image (mean ± SD). To detect cortical layers 1–6 (middle), three fixed points (●) and one reference point (▪) were selected; for layers 1–5 (right), the same three points and one alternative point (★) were used. Representative dendrites are indicated by arrows and arrowheads. Scale bar in raw data image, 200 µm. Orientation angles were defined with respect to the dorsal–ventral (D–V) axis, where the ventral direction was set as 0 degrees. D: dorsal; V: ventral; M: medial. (D) CTIP2 staining for visualization and transformation of layers 5 and 6. Point selection followed the same convention as in (C). Scale bar in raw data image, 200 µm. (E) Quantification of the dorsal–ventral differences in the starting position of layer 5 after transformation. mean ± SD. *n* = 12. Paired *t*-test. *** *p* < 0.001; n.s. not significant.

To accurately quantify synaptic organization along the depth axis of the ACC, we referred to anatomical coordinates based on the Allen Brain Atlas (Allen Institute for Brain, 2011). Initially, we applied a perspective transformation to straighten the cortical curvature, which allowed for approximate alignment along the depth axis. However, this process disproportionately stretched the ventral region, risking overrepresentation of ventral signals when averaging data. To address this distortion, we implemented an additional logarithmic transformation following the perspective transformation, aiming to equilibrate dorsal-ventral contributions. Optimization of the logarithmic constant revealed that a coefficient of a = 0.015 best preserved the uniformity across the depth axis (Figure 2B and Figure 2—figure supplement 1B).

To validate the effectiveness of this transformation, we first constructed an artificial geometric model simulating dendritic orientation. Application of our pipeline to this model confirmed both isotropy (uniform orientation) and equidistant spacing across the normalized depth axis (Figure 2—figure supplement 1C). To assess the reproducibility of line drawing performance, we used actual immunostained images and quantified spacing and slope consistency across 10 repeated trials. One-way ANOVA and Levene’s test revealed no significant differences in interline spacing (ANOVA *p* = 0.4310; Levene *p* = 0.2119) or slope direction (ANOVA *p* = 0.1309; Levene *p* = 1.0000) across trials, indicating robust inter-trial consistency in both spacing and directionality (Figure 2—figure supplement 1D and E).

We next tested whether the transformation improved the isotropy of actual dendritic structures in the ACC. Applying the transformation to sections stained for MAP2, a dendritic marker, we observed that the average dendritic orientation was corrected from 30.7° ± 40.5° before transformation to 2.2° ± 53.1° after transformation relative to the medial-lateral axis (Figure 2C left and middle).

Finally, we examined how this transformation affected layer-specific structure. In the artificial model, deeper cortical layers (L5 and L6) appeared at shallower depths dorsally than ventrally (Figure 2B). Specifically, the L1–L2/3 boundary appeared at 5.4% dorsally and 21.2% ventrally, the L2/3–L5 boundary at 14.5% dorsally and 52.6% ventrally, and the L5–L6 boundary at 31.4% dorsally and 90.8% ventrally (Figure 2—figure supplement 1F). Consistent with this, immunostaining for CTIP2—a molecular marker of deep cortical layers—demonstrated that L5 onset occurred at 11.72% ± 2.55 dorsally and 41.22% ± 8.99 ventrally (Figure 2D middle and E). These results confirm that, without correction, the original depth transformation introduced a dorsal-ventral misalignment of layer boundaries, particularly in the deep layer. To further correct for this discrepancy, we refined the transformation by excluding L6 from the depth normalization and adjusted the logarithmic coefficient to a = 0.002. Under these conditions, both dendritic isotropy and the dorsal-ventral alignment of L5 onset were successfully achieved (Figure 2C right, 2D right and 2E).

These results collectively validate our transformation pipeline as an effective method for minimizing regional distortion and enabling accurate depth-specific quantification of synaptic organization in the ACC.

### Reduced inhibitory synapse density leads to elevated structural E/I ratio in the ACC of VPA-exposed mice

We next applied the transformation pipeline to quantify excitatory and inhibitory synaptic organization along the depth axis of the ACC. Immunostaining for PSD-95 and gephyrin was performed to visualize excitatory and inhibitory synapses, respectively (Figure 3A). Following perspective and logarithmic transformation, the ACC was divided into tiles, and synaptic puncta were automatically counted in each tile. To minimize misdetection at the boundary between layers 5 and 6, we employed the L1–6 version of the transformation pipeline. Synaptic densities and area were then plotted along the normalized depth axis, approximating cortical layer boundaries.

**Figure 3.**
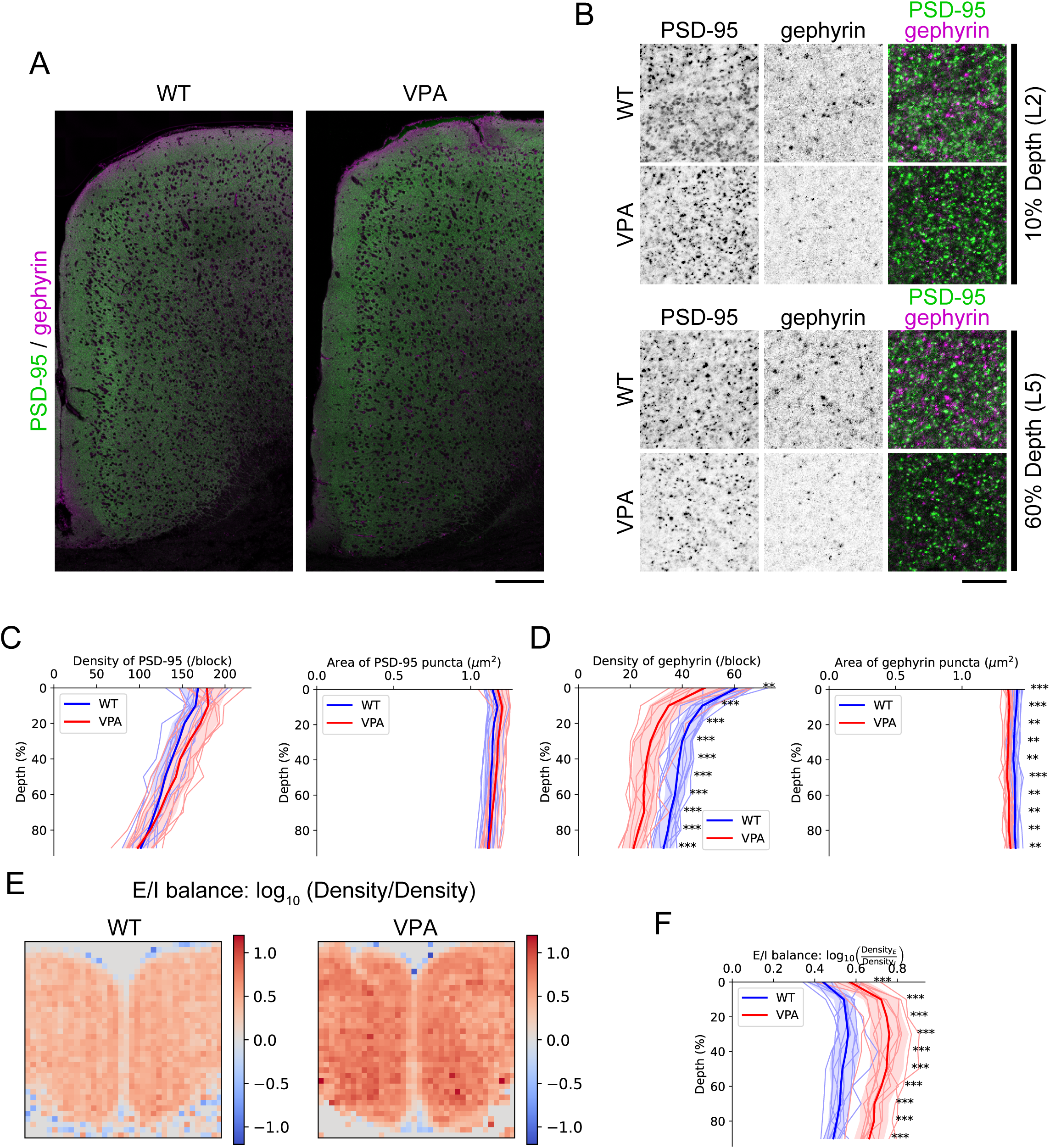
Analysis of synaptic markers and E/I balance in VPA model mice using the standardized pipeline. (A) Immunohistochemical visualization of PSD-95 and gephyrin across the entire ACC. Scale bar, 200 µm. (B) Enlarged views of representative regions at 10% depth (corresponding to layer 2/3) and 60% depth (corresponding to layer 5). Scale bar, 10 µm. (C) Depth-dependent distribution of PSD-95 density and area. Each block represents a 53.4 µm × 53.4 µm region. Thick lines indicate group means; thin lines represent individual values; shaded areas denote mean ± SD. (D) Depth-dependent distribution of gephyrin density and area. Block size and graphical conventions are the same as in (C). (E) Reconstructed E/I balance map. The balance was calculated from quantitative values within each block. (F) Depth-dependent profiles of E/I balance. For (C), (D), and (F): WT (*n* = 11), VPA (*n* = 11) Welch’s *t*-test. ** *q* < 0.01; *** *q* < 0.001 after BH FDR. n.s. (not significant) is not indicated.

In both WT and VPA-exposed mice, excitatory and inhibitory synapse densities exhibited characteristic laminar profiles, with peaks observed in the superficial layers. Specifically, VPA-exposed mice exhibited a trend toward increased excitatory synapse density in the superficial layers compared to WT controls (e.g., density at 10% depth: WT = 165.4 ± 9.9, VPA = 180.2 ± 13.5, *q* = 0.081 after Benjamini–Hochberg (BH) correction), whereas no significant difference was observed in synapse area (e.g., area at 10% depth: WT = 1.15 ± 0.043 µm^2^, VPA = 1.19 ± 0.039 µm^2^, *q* = 0.25) (Figures 3B and C). For inhibitory synapses, both the density and area of gephyrin-positive puncta were elevated across cortical depths in VPA-exposed mice (e.g., density at 10% depth: WT = 47.7 ± 4.7, VPA = 34.7 ± 7.3, *q* = 0.0001; area at 10% depth: WT = 1.40 ± 0.020 µm^2^, VPA = 1.36 ± 0.027 µm^2^, *q* = 0.0004) (Figures 3B and D).

To further assess the balance between excitatory and inhibitory inputs, we calculated the E/I balance across cortical depths. The reconstructed anatomical maps and depth-wise graphs demonstrated a significant increase in the E/I balance throughout the cortical depth in VPA-exposed mice, with a tendency toward a more pronounced effect in the ventral deep layers (Figures 3E and F).

Finally, we performed qPCR analysis on laser-microdissected ACC samples to measure Dlg4 (commonly known as PSD-95) as an excitatory synapse marker and Nlgn2 (Neuroligin-2) as an inhibitory synapse marker (Figure 3—figure supplement 1A) (Poulopoulos et al., 2009); however, both genes exhibited substantial variability, and no significant difference was observed between groups (Figure 3—figure supplement 1B). Although partial RNA degradation was observed (Figure 3—figure supplement 1C and D), the overall quality was sufficient for analysis, suggesting that the variability is more likely attributable to individual differences.

These findings suggest that prenatal VPA exposure induces a depth-dependent shift in excitatory and inhibitory synaptic organization within the ACC, leading to an overall increase in the E/I balance and potential alterations in cortical circuit function.

### Multivariate analysis integrating behavioral and synaptic parameters

To investigate how prenatal VPA exposure affects behavior and synaptic organization, we first computed pairwise Pearson correlations between behavioral scores and synaptic measures across five cortical depth bins (0–10%, 10–20%, 20–40%, 40–70%, 70–100%) based on cortical laminar architecture (Figure 2E and Figure 2—figure supplement 1F). Because the E/I ratio is highly sensitive to variance in inhibitory synapses (Figure 3D and F) and lacks a well-defined linear biological correspondence (Baker et al., 2020; Bhatia et al., 2019; Miehl & Gjorgjieva, 2022), we analyzed excitatory (PSD-95) and inhibitory (gephyrin) synaptic densities separately.

To capture the uncorrected exploratory trend, we first analyzed the data without correction. Excitatory synapses showed correlations with working memory in the superficial to middle layers, whereas inhibitory synapses were correlated with Y-maze re-entry rate and social interaction in the middle to deep layers (Figure 4—figure supplement 1A). Next, to correct for false positives due to multiple comparisons, we applied the BH False Discovery Rate (FDR) correction (α = 0.05) to pre-specified comparisons in the superficial layer (10–20%), based on our primary hypothesis derived from the histological results (Figure 3C and D). PSD-95 density showed significant correlations with Y-maze performance (alternation rate: *r* = −0.57, *q* = 0.031; re-entry ratio: *r* = 0.65, *q* = 0.0135), whereas gephyrin showed no significant associations (Appendix 1—table 1). When correcting for correlations across all 60 combinations (6 behaviors × 2 markers × 5 bins), no comparisons remained significant after FDR correction (Appendix 1—table 2).

Although none of the 60 individual correlations survived FDR adjustment, this stringent control may inflate false-negative rates when variables are highly inter-correlated. We therefore applied a multivariate strategy (PCA) to capture shared variance among synaptic and behavioral measures while drastically reducing the multiplicity burden. PCA revealed clear separation between WT and VPA-exposed mice along the first two principal components (PCs), with minor overlap (Figure 4A). Focusing on the top three behavioral indicators based on PC1 and PC2 loadings (Figure 4B), we examined the results of correlation analysis. Density of PSD-95, Y-maze alternation rate, and re-entry ratio were all strongly correlated with PC1 in either the positive or negative direction (*r* = +0.76, −0.80, and +0.86, respectively). Gephyrin density was also negatively correlated with PC1 (*r* = −0.62). In contrast, open-field center time and elevated plus-maze open-arm time showed strong negative correlations with PC2 (*r* = −0.87 and −0.75, respectively) (Figure 4C). These loadings supported the interpretation of PC1 as reflecting ACC-related excitability and PC2 as indexing anxiety-related behavioral traits (Figure 4—figure supplement 2A).

**Figure 4.**
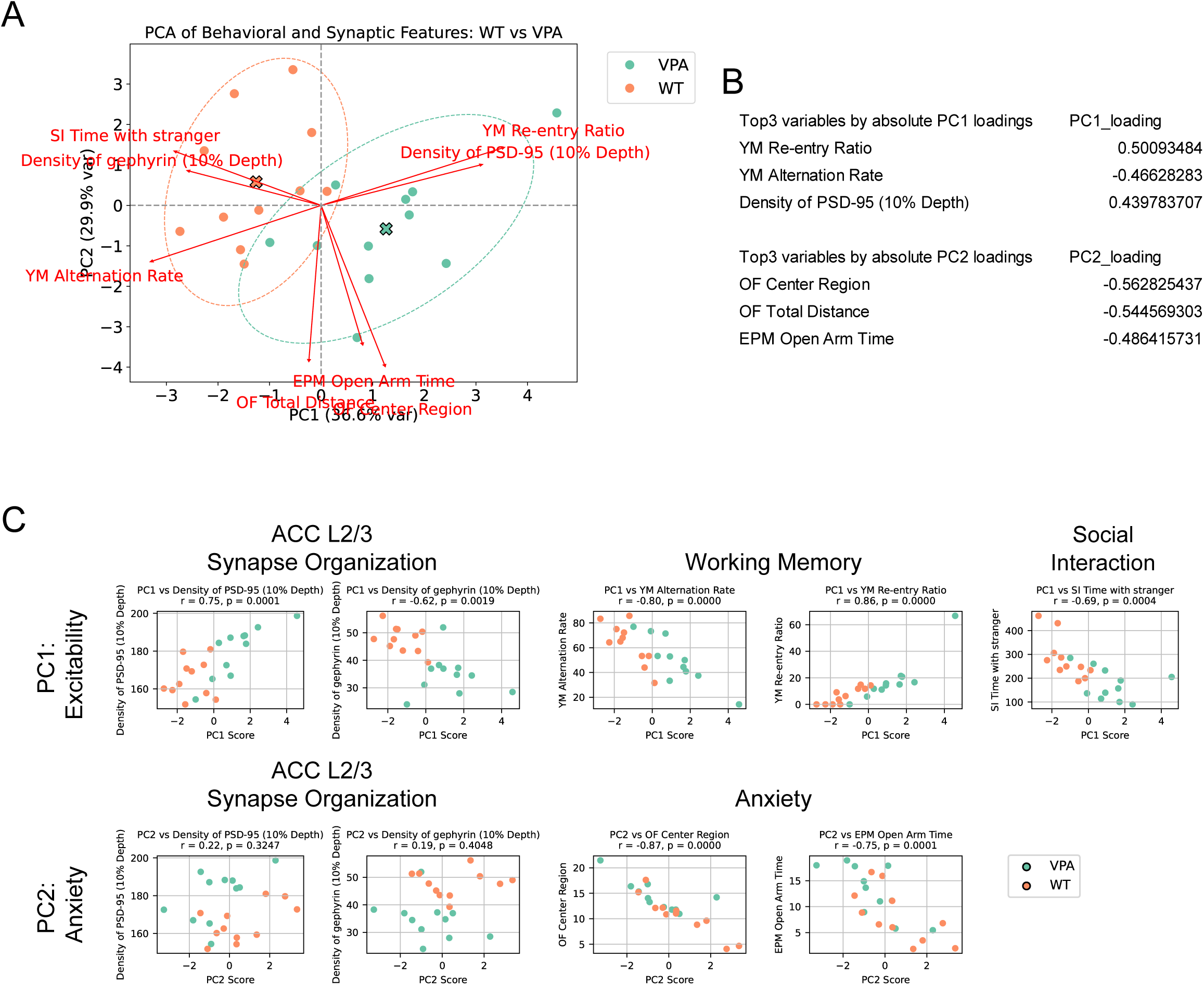
Multivariate analysis of synaptic and behavioral features in WT and VPA model mice. (A) PCA constructed from synaptic organization and behavioral data, showing clear group separation between WT and VPA mice. The plot includes a vector diagram indicating the contribution of each variable to the PCA components. Centroids are indicated by × marks, and arrows denote their displacement. The dashed ellipses represent 95% confidence intervals (CIs) for each group.PC1 and PC2 explained 36.6% and 29.9% of the total variance, respectively. (B) Top three variables contributing most strongly to PC1 and PC2, ranked by the absolute value of their loadings. (C) Correlation plots between representative original variables and the principal components (PC1 and PC2). WT (*n* = 11), VPA (*n* = 11). These results suggest that PC1 reflects excitability, while PC2 is associated with anxiety-related traits.

To evaluate the robustness of our PCA-based feature space, we applied multiple validation approaches. Bootstrap PCA (1,000 iterations) confirmed stable loadings and explained variance across all components (Figure 4—figure supplement 2B and C). PC3 and PC4 did not contribute to group separation and were excluded from further analysis (Figure 4—figure supplement 2D). To assess classification performance in PCA space, we trained logistic regression and SVM classifiers using the eight PCA components. Both models showed high accuracy and perfect AUCs, with permutation testing (*p* = 0.001) confirming that group separation was highly unlikely to occur by chance (Appendix 2—table 1). Additionally, five-fold cross-validation of PCA-based clustering (Figure 4—figure supplement 2E) demonstrated high compression efficiency, further supporting the robustness of the PCA-defined feature space.

As a complementary approach, we also explored an alternative multivariate analysis method. To assess group discriminability, we applied linear discriminant analysis (LDA), which achieved maximal separation between WT and VPA-exposed mice (Figure 4—figure supplement 2F and G). Contribution analysis indicated that density of PSD-95 and open-field center time were major drivers. However, social interaction, which represents a core behavioral impairment in VPA-exposed mice, contributed minimally and exhibited inconsistency across the three indices (Appendix 3—table 1). These discrepancies led us to compare LDA and PCA directly, and PCA more consistently captured the phenotypic divergence observed in our pipeline.

Collectively, these findings highlight that PCA played a central role in distinguishing VPA-exposed mice from WT, integrating individual behavioral and synaptic variables into a coherent classification framework. This supports the notion that prenatal VPA exposure induces convergent disruptions across behavior and neural circuits, and demonstrates the utility of multivariate profiling for capturing biologically meaningful, high-dimensional phenotypes.

### Spontaneous resilience within the VPA group

To evaluate the broader applicability of our analysis pipeline, we proceeded to investigate individual variability and potential intervention effects. Building on the robust group separation observed in PCA space (as demonstrated in the previous section), we next focused on individual differences within the VPA-exposed group. Behavioral tests, immunohistochemistry, and qPCR all revealed considerable within-group variability in the VPA mice, making group-level interpretation challenging (Figure 1, Figure 3, and Figure 3—figure supplement 1). To better capture this individual variability, we classified VPA mice based on their *Z*-distance from the WT centroid in PCA space (Figure 5A). Mice with *|Z|* ≤ 1.96 were operationally defined as “resilient” (*n* = 2). Given the small sample size, this analysis is exploratory in nature and intended to generate hypotheses.

**Figure 5.**
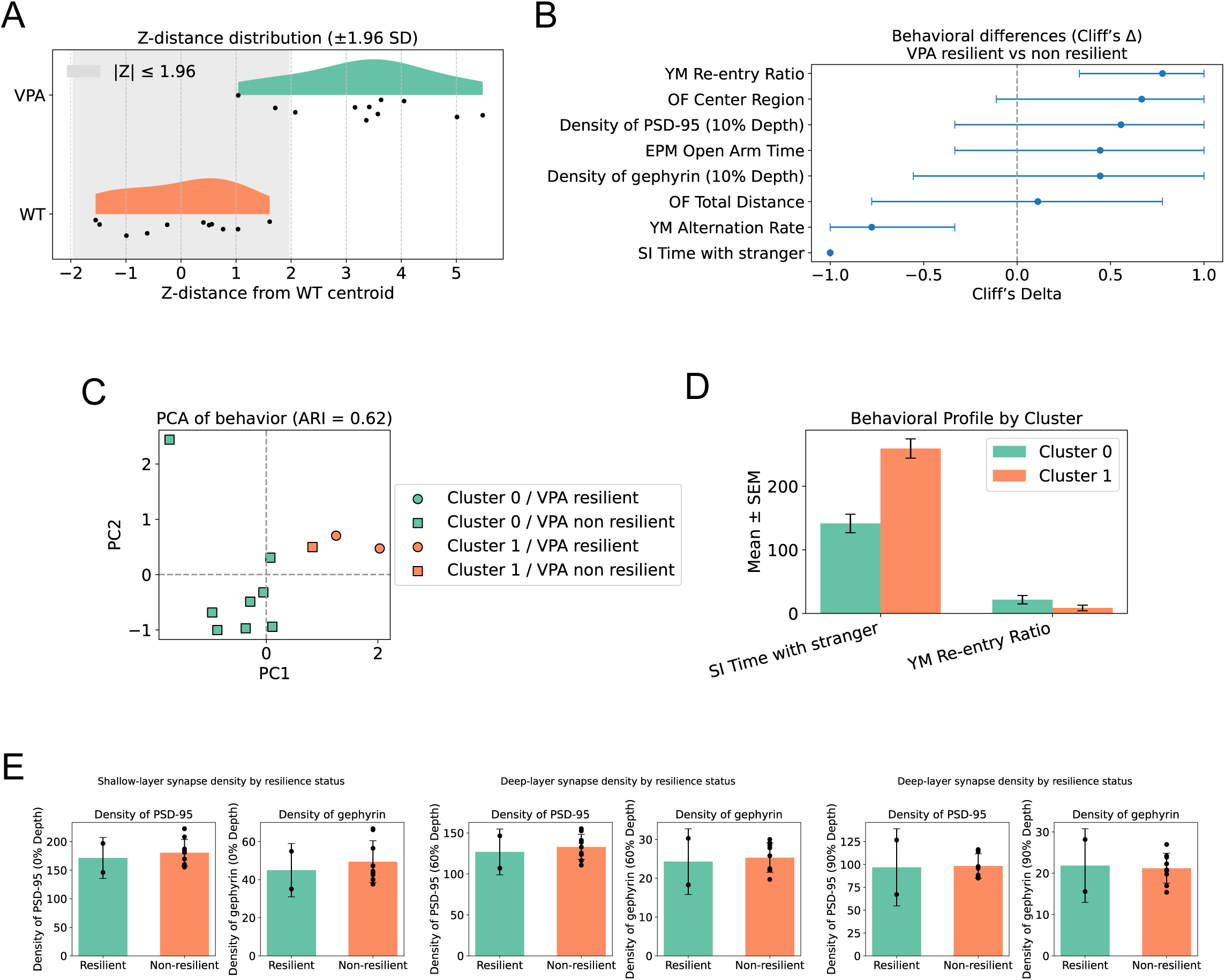
Identification and characterization of resilient individuals within the VPA group. (A) Raincloud plot showing Z-distance from the WT group centroid. Two VPA-exposed individuals fell within 1.96 standard deviations and were classified as resilient. WT (*n* = 11), VPA (*n* = 11). (B) Cliff’s Δ analysis comparing resilient and non-resilient individuals within the VPA group. Three behavioral variables—Y-maze re-entry ratio, alternation rate, and social interaction time with a stranger—had 95% confidence intervals that did not cross zero, indicating meaningful separation between subgroups. (C) Adjusted Rand Index (ARI) analysis based on k-means clustering using two ASD-related behavioral phenotypes (social interaction time and Y-maze re-entry ratio). The resulting ARI score was 0.62, and the cluster structure successfully separated the two non-resilient individuals. *n* = 11. (D) Comparison of behavioral performance by cluster (mean ± SEM). Significant differences were observed in social interaction time and Y-maze re-entry ratio between clusters. Cluster 0 (*n* = 8), Cluster 1 (*n* = 3). (E) Synaptic organization comparison between resilient and non-resilient VPA mice. Although the analysis focused on cortical depths not used in the clustering, no significant differences were found. mean ± SD. Resilient (*n* = 2), Non-resilient (*n* = 9). Since *n* < 3, statistical significance testing was not performed.

Effect size analysis (Cliff’s *Δ*) showed that resilient mice had longer social interaction time, higher alternation rate, and lower Y-maze re-entry ratio, with 95% confidence intervals not crossing zero (Figure 5B). Unsupervised clustering using two behavioral features—social interaction time and re-entry ratio—closely matched the *Z*-based classification (ARI = 0.62), suggesting a distinct behavioral profile (Figure 5C). Including alternation rate reduced classification performance, indicating feature specificity. Although leave-one-out validation was not feasible due to sample size, PCA proximity and consistent behavioral patterns support the classification of these two mice as a distinct subgroup (Figure 5C and D).

Finally, we explored whether resilience was reflected in synaptic phenotypes not used in the *Z*-score calculation. Specifically, we examined synaptic densities of PSD-95 and gephyrin in the remaining cortical layers (0%, 60%, and 90% depth). Although the resilient group comprised only two mice, exploratory bar plots did not reveal any notable group differences (Figure 5D). These findings suggest that resilience may not be fully explained by region-wide, depth-dependent synaptic organization within the ACC alone, indicating the potential need for finer subdivisions—such as along the dorsal–ventral axis or between local and long-range circuits. Given the limited sample size, these results should be considered preliminary and hypothesis-generating

### Exploratory analysis of voluntary exercise effects on synaptic and behavioral phenotypes

To assess whether our synaptic–behavioral pipeline is sensitive enough to detect intervention effects, we conducted an exploratory study involving voluntary wheel running in VPA-exposed mice (*n* = 3 per group) (Figure 5—figure supplement 1A). Behaviorally, exercise enhanced social interaction (*p* = 0.049, Hedges’ *g* = 1.22) and reduced perseverative re-entries (*p* = 0.046, *g* = −0.98), though effects on anxiety-related measures and locomotion were inconsistent (Figure 5—figure supplement 1B).

Synaptically, exercise increased both excitatory and inhibitory synapse densities, particularly at 90% cortical depth (excitatory: *q* = 0.0017; inhibitory: *q* = 0.020 after BH), resulting in a shift in E/I balance toward WT levels (Figure 5—figure supplement 1C and D). When projected into the PCA space defined by control animals, WT mice with exercise appeared separated from the control WT cluster, whereas VPA mice with exercise remained within the distribution of the control group but shifted toward the WT control and WT with exercise clusters (Figure 5—figure supplement 1E). To verify that the intervention did not introduce an entirely new dominant axis of variance, we recomputed the PCA including all animals. The cumulative variance explained by the first two principal components remained comparable (66.5% → 59.8%; Figure 5—figure supplement 1F), and Procrustes analysis further confirmed that the overall structure of the PCA space was preserved (similarity = 0.96; cosine similarity for PC1 = 0.992, PC2 = 0.871). These results indicate that exercise induced a translation within the existing feature space rather than introducing a qualitatively distinct dimension of variance. WT mice with exercise showed a significant multivariate shift (Hotelling’s *T*² = 38.6, *p* = 0.0004), while VPA mice with exercise exhibited a smaller, non-significant displacement (Appendix 4—table 1 and 2). A similar trend was observed in the bootstrap analysis (Figure 5—figure supplement 1G). However, given the limited sample size, all results should be interpreted as preliminary and hypothesis-generating. Future studies with ≥19 mice per group, ideally powered to detect moderate effects (Figure 5—figure supplement 1H), will be essential to confirm the partial phenotypic rescue observed here and to elucidate the underlying circuit mechanisms.

## Discussion

In this study, we developed a standardized analysis pipeline combining perspective transformation, logarithmic scaling, and multivariate profiling, enabling depth-aligned, quantitative mapping of excitatory and inhibitory synaptic organization alongside behavioral assessments. By applying this pipeline to a prenatal VPA exposure model of ASD, we identified a trend toward depth-specific disruptions in excitatory synaptic organization, alterations in inhibitory synapse density across cortical layers, and associations with core ASD phenotypes such as impaired social behavior and working memory. Furthermore, although exploratory, the robustness of PCA enabled identification of resilient individuals and evaluation of potential intervention effects. Together, our findings highlight the utility of integrated synaptic and behavioral profiling for dissecting circuit-level abnormalities and assessing therapeutic outcomes in neurodevelopmental disorders.

### Circuit-level disruptions in the VPA model

VPA-exposed mice exhibited impaired social interaction, consistent with previous reports (Nicolini & Fahnestock, 2018), but showed reduced anxiety-like behaviors—contrary to many prior studies (Mabunga et al., 2015; Mehta et al., 2011). This discrepancy may reflect methodological differences, including VPA administration protocol and testing environment. Despite the lack of statistically significant group-level differences in repetitive behavior (Figure 1C), PCA focused on superficial ACC layers revealed a trend toward increased excitatory synapse number, particularly at 10% normalized depth corresponding to L2/3 (Figure 3C), was a major contributor to PC1 (loading = 0.440), which correlated with elevated Y-maze re-entry ratio (Figure 4). These results suggest that excessive superficial excitatory input may underlie cognitive deficits, including impaired working memory, though the possibility of stereotypy must also be considered.

Although gephyrin density in deeper layers (70-100% depth) moderately correlated with several behavioral readouts (e.g., Y-maze re-entry ratio: *r* = −0.52; social-interaction time: *r* = 0.47; EPM Open arm time: *r* = −0.39), PCA restricted to superficial data (10 % depth) was dominated by excitatory variables (PC1), showing weaker associations with anxiety (PC2). Because these two analyses probe different cortical depths, the apparent discrepancy likely reflects depth-specific circuit properties. This raises the possibility among other factors that variations in interneuron subtype composition along the depth axis (e.g., parvalbumin [PV]-, somatostatin [SST]-, and vasoactive intestinal peptide [VIP]-expressing neurons) contribute to behaviorally relevant inhibitory signaling (Shao et al., 2021; Tremblay et al., 2016). Future studies employing subtype-specific markers will be needed to clarify this point. Although group means of Neuroligin-2 were similar, its high inter-individual variability suggests potential differences in inhibitory circuit maturation, warranting further investigation (Figure 3—figure supplement 1B).

Correlations between anxiety-related metrics (e.g., open field center time, EPM open arm time) and PCA axes were modest (*r* = 0.30 and 0.19 for PC1, respectively), suggesting limited association with L2/3 excitatory synaptic changes. Notably, we did not assess excitatory amygdala-to-ACC inputs or distinguish synaptic contacts onto pyramidal neurons versus interneurons (Lee et al., 2021) nor did we examine synaptic alterations within the amygdala itself (Lin et al., 2013). These limitations highlight the need for circuit- and cell-type-resolved approaches. Future work may involve manipulation of specific ACC–amygdala pathways to test whether depth-specific synapse formation is altered and how distinct behavioral parameters shift, using the multiscale pipeline developed here.

Although exploratory in nature, voluntary exercise appeared to partially rescue both synaptic and behavioral phenotypes in VPA-exposed mice (Figure 5—figure supplement 1). Significant increases in synaptic density were observed in deep ACC layers, which receive input from midline thalamic nuclei involved in cognitive and motivational processing (Ito et al., 2015; Vertes, 2006; Vertes et al., 2022). These circuit-level changes coincided with improvements in working memory, consistent with previous reports (Wan et al., 2024). Furthermore, both depth-resolved quantification and PCA analysis indicated that exercise intervention shifted the E/I profile of VPA-exposed mice closer to that of WT controls.

Interestingly, exercise also induced a significant shift in WT mice along the positive direction of PC1 (excitability) and the negative direction of PC2 (anxiety), which was unexpected (Hotelling’s *T*² = 38.6, *p* = 0.0004, and Figure 5—figure supplement 1G). This suggests that even modest environmental interventions can modulate synaptic and behavioral states within the ACC. Consequently, the PCA space that WT animals can occupy may be broader than observed under standard housing conditions. If so, ASD-like phenotypes may be more accurately confined to the upper-right quadrant of PCA space (high PC1, high PC2). Taken together with the results of our resilience classification analysis, this interpretation suggests that a considerable proportion of VPA-exposed mice exhibit a resilient phenotype (Figure 5—figure supplement 1I). In future work, these possibilities should be tested in larger cohorts to assess the robustness and generalizability of our findings.

Together, these findings suggest that layer-specific E/I imbalance—particularly in deep circuits—can be modified by experience-dependent plasticity. Clarifying how this plasticity engages dynamic intracellular signaling pathways related with synaptic organizers (Ichinose et al., 2015; Ichinose et al., 2023) will be a key step toward understanding E/I balance regulation and ASD pathophysiology. Furthermore, such knowledge will offer a valuable point of contrast to gene-defined models, such as Shank3-deficient mice (Peça et al., 2011), enabling comparative analysis of circuit-level pathology arising from environmental versus genetic origins.

### Development and Application of a Cross-Level Analysis Pipeline

Our analysis pipeline addresses a critical methodological gap: quantitatively linking synaptic architecture to behavioral phenotypes across cortical structures. While approaches like Brainbow have enabled detailed morphometric analyses, and classical methods such as Golgi staining have elucidated neuronal morphology (Megías et al., 2001; Yoshihara et al., 2021), Golgi staining faces depth-related limitations and lacks reliable resolution of inhibitory synapses (Ranjan & Mallick, 2010; Spruston, 2008).

Traditional immunohistochemistry also involves complex protocols, often requiring antigen retrieval for adequate labeling. In contrast, our pipeline—combining glyoxal-based immersion fixation, vibratome sectioning, and standard antibody labeling—offers a more accessible and efficient approach (Konno et al., 2023). High-resolution tile imaging further enables rapid acquisition of large-scale synaptic datasets (Balasubramanian et al., 2023).

To ensure depth-wise accuracy, we implemented perspective transformation and logarithmic scaling, minimizing tissue distortion and depth bias. This allows for depth-aligned, uniform synaptic mapping and holds promise for large-scale screening applications. Given outstanding challenges in analyzing the basolateral amygdala (BLA) in the VPA model, we aim to extend this pipeline to the BLA and eventually toward whole-brain mapping.

Despite achieving depth normalization, resolving dorsoventral spatial patterns remains difficult. Although E/I balance maps were successfully generated for individual animals, anatomical variability still hampers group-level comparisons. As a next step, we will implement AI-based morphological standardization (Carey et al., 2023; Sadeghi et al., 2023) enabling consistent 2D and 3D statistical analyses across individuals and regions.

By integrating multivariate profiling, our pipeline successfully distinguished VPA-exposed mice from WT controls based on convergent structural and behavioral abnormalities. This robustness also enabled the identification of resilient individuals within the VPA group. Although exploratory (*n* = 3 per exercise group), the data further suggested that voluntary exercise partially restored synaptic E/I balance and improved social behavior. These findings demonstrate the utility of our pipeline for both mechanistic studies and preclinical evaluation of therapeutic interventions in ASD.

To more fully bridge synaptic architecture and behavior, incorporating intermediate physiological readouts will be essential. *In vivo* electrophysiology or calcium imaging could capture dynamic circuit activity, while noninvasive techniques like resting-state fMRI or functional ultrasound may support cross-scale, multimodal integration. Furthermore, although this study could not apply such integration due to differing fixation protocols between immunohistochemistry and qPCR, combining synaptic mapping with spatial transcriptomics or single-cell analysis is expected to enable identification of molecular features associated with circuit disruptions and behavioral phenotypes.

### Technical limitations and why univariate correlations remain weak

Quantitative synaptic imaging faces challenges from both sampling error and technical variability. In our 80 µm sections, minor warping during mounting caused local z-depth shifts, and uneven antibody penetration (e.g., PSD-95 labeling was predominantly restricted to superficial z-planes, in contrast to deeper gephyrin signals) introduced depth-dependent intensity loss. While histogram matching corrected global brightness and contrast, fine-scale artefacts—such as z-attenuation, tile-edge loss, and lens aberrations—persisted as spatially irregular, channel-specific noise.

Such noise undermines univariate analyses: single-marker fluctuations may reflect technical issues rather than biology, obscuring true behavior–circuit links. Compounding this, behaviors like social interaction or working memory emerge from distributed circuit activity, not from synapse counts at a single depth. Additionally, each synaptic marker encompasses heterogeneous sources (e.g., PV-, SST-, and VIP-interneurons for gephyrin), so changes in puncta number may not map linearly onto circuit function. If circuit-level effects are nonlinear or involve multiple co-modulated markers, single-variable correlations underrepresent biological relevance.

To overcome these limitations, we adopted a two-step approach: histogram matching reduced baseline intensity disparities, and PCA extracted biologically relevant co-variation across markers. For instance, PC1 captured coordinated changes in PSD-95, gephyrin, and E/I balance aligned with behavioral variation, while unstructured noise was relegated to higher-order PCs. PCA thus uncovered circuit– behavior links missed by univariate methods.

Future refinements—such as z-stack calibration or voxel-wise attenuation correction—may further suppress artefacts. Still, our pipeline illustrates a key principle: multivariate analysis can recover meaningful structure–function relationships obscured by variable-specific noise.

In summary, this study provides a novel cross-level analytical framework linking cortical synaptic organization to behavioral phenotypes, and identifies depth-specific disruptions of E/I balance in a widely used ASD model. Our pipeline successfully captured both structural and behavioral abnormalities, and demonstrated potential utility for identifying resilient individuals as well as detecting the effects of therapeutic interventions. By addressing key methodological gaps, this approach opens avenues for more precise profiling of circuit-level pathology and behavioral outcomes. Future expansions toward broader brain regions and integration with functional modalities will further enhance its applicability in neurodevelopmental disorder research.

## Materials & methods

### Animals and VPA-induced ASD model mice

C57BL/6J mice (JAX strain; Jackson Laboratory Japan; IMSR Cat# JAX:000664, RRID:IMSR_JAX:000664) were bred in-house at the Gunma University animal facility. For timed pregnancy, one male and two females were paired in the late afternoon, and the presence of a vaginal plug was checked the following morning. The day a plug was detected was designated as embryonic day 0.5 (E0.5). Pregnancy was further confirmed by monitoring body weight gain, and only females that exhibited appropriate weight gain by embryonic day 12.5 (E12.5) were used for VPA administration. On E12.5, pregnant dams were intraperitoneally injected with VPA (500 mg/kg, sodium valproate, FUJIFILM-Wako, Japan; Cat# 193-18352) dissolved in sterile PBS. Control dams received an equivalent volume of PBS and did not result in any observable adverse effects. All offspring were weaned at postnatal day 21 (P21) and housed in same-sex cages under a 12-hour light/dark cycle with ad libitum access to food and water. To avoid isolation-induced stress, animals were never singly housed and were instead group-housed with littermates whenever possible, in 225 × 338 × 140 mm transparent polycarbonate cages (CLEA Japan, Inc., Japan). Behavioral tests were conducted at postnatal 12 weeks, and brain tissue was collected soon after the behavioral tests for histological and molecular analysis. All animal procedures were approved by Gunma University for animal experiments (approval number: 24-056), in accordance with institutional and national guidelines.

### Behavioral analyses

Behavioral analyses were conducted during the light phase in a quiet, controlled environment using male mice at 12 weeks of age. Mice were habituated to the testing room for at least 30 minutes prior to each test. Behavioral experiments were performed by experimenters who were aware of group assignments. However, all behavioral metrics were quantified using automated tracking systems and predefined objective criteria to minimize observer bias. Heatmaps of dwell time were generated by averaging binarized images in ImageJ, and movement trajectories were automatically computed using the manufacturer’s analysis software.

### Social interaction test (modified open field-based)

To evaluate social behavior, we employed a modified open field paradigm instead of the standard three-chamber test. A acrylic open field arena (50 × 50 × 30 cm, O’Hara & Co., Ltd, Japan) was used, with two quarter-circle wire mesh cages (diameter: 20 cm) placed in diagonally opposite corners (Figure 1A). The subject mouse was habituated to the arena for 10 minutes without the target mouse. During the test session, a stranger mouse (sex- and age-matched) was placed in the cage located in the lower left corner, while the cage in the upper right corner remained empty (Figure 1A). The time spent in the interaction zone (a 6-cm-wide area around the cage) and the number and time of entries into this zone were recorded as indices of social interest. The visit ratio to the stranger was calculated as the number of visits to the stranger mouse divided by the total number of visits to both cages. Data were analyzed using the manufacturer’s software.

### Y-maze spontaneous alternation

Spontaneous alternation behavior was assessed using a Y-maze composed of three arms (40 × 3 × 12 cm, 120° angle between arms, O’Hara & Co., Ltd.) (Figure 1A). Mice were placed at the end of the arm and allowed to explore freely for 10 minutes. The sequence of arm entries was recorded with the manufacturer’s software, and spontaneous alternation (%) was calculated as the number of triads consisting of entries into all three different arms divided by the total number of possible alternations. In addition to alternation, the total number of arm entries and re-entries (repeated entries into the same arm within three consecutive entries) were also measured as indicators of repetitive behavior.

### Elevated plus maze (EPM)

The elevated plus maze (EPM) apparatus consisted of two open arms (30 × 5 cm) and two closed arms (30 × 5 × 20 cm) arranged in a cross and elevated 50 cm above the floor (Figure 1A). Each mouse was placed at the end of an open arm and allowed to explore freely for 10 minutes. Time spent in the open arms was recorded. The video data were edited into 300 × 300 pixel movies at 6 frames per second, and analysis was performed using a custom analysis pipeline based on ImageJ and Python.

### Open field test

Locomotor activity and anxiety-like behavior were evaluated in the same open field arena (Figure 1A). Mice were placed in the center and allowed to explore for 10 minutes. Total distance traveled and time spent in the center zone (30 × 30 cm) were analyzed using the manufacturer’s software.

### Voluntary exercise

To provide voluntary exercise, mice were housed in transparent polycarbonate cages equipped with a low-resistance running wheel (Fast-Trac, Bio-Serv, #K3250) (Figure 5—figure supplement 1A). The wheel was freely rotating and accessible throughout the intervention period. Bedding and food were provided ad libitum, and animals were allowed to engage in voluntary running without any external stimuli or constraints.

### Immunohistochemistry following glyoxal fixation

Following a previously reported protocol (Konno et al., 2023) with modifications, mice were euthanized by cervical dislocation, and brains were rapidly removed and immersed in a fixative solution containing 4% glyoxal and 9% acetic acid, adjusted to pH 4.5 using 1 M NaOH, for 40 minutes at room temperature. For MAP2 immunostaining, however, brains were fixed overnight in 4% paraformaldehyde (PFA) in PBS at 4 °C. After two washes with PBS, coronal brain sections (80 μm thick) were cut using a vibratome (Leica VT1000 S). ACC regions were sampled from coronal sections centered around the anatomical landmark where the left and right corpus callosum converge, corresponding approximately to Bregma +1.2 mm (±0.24 mm). Sampling was guided by the anatomical features rather than absolute stereotaxic coordinates to account for individual variability. Free-floating sections were permeabilized with 2% Triton X-100 in PBS for 30 minutes and blocked in 2% normal donkey serum for 30 minutes at room temperature. Sections were then incubated overnight at room temperature with the following primary antibodies: rabbit anti-PSD-95 (1:1000, Cell Signaling Technology, Cat# 36233, RRID:AB_2721262), mouse anti-gephyrin (1:1000, Synaptic Systems, Cat# 147011, RRID:AB_887717), rat polyclonal anti-CTIP2 (1:1000, Abcam, Cat# ab18465, RRID:AB_2064130) and chicken anti-MAP2 (1:10000, Novus, Cat# NB300-213, RRID:AB_2138178). After thorough washing, sections were incubated for 1 hour at room temperature with fluorophore-conjugated secondary antibodies: donkey anti-rat IgG (H+L) Alexa Fluor 488 (1:1000, Jackson ImmunoResearch, Cat# 712-546-150, RRID:AB_2340685), donkey anti-rabbit IgG (H+L) CF568 (1:2000, Biotium, Cat# 20098-1, RRID:AB_10853318), donkey anti-mouse IgG (H+L) CF633 (1:2000, Biotium, Cat# 20124, RRID:AB_10557033), and donkey anti-chicken Chicken IgY (IgG) (H+L), Alexa Fluor 647 (1:1000, Jackson ImmunoResearch, Cat# 703-605-155, RRID:AB_2340379). Following immunostaining, sections were post-fixed in 4% paraformaldehyde (PFA) in PBS for 10 minutes. Finally, sections were mounted on glass slides and coverslipped using ProLong™ Glass Antifade Mountant with NucBlue Stain (Thermo Fisher Scientific, Cat# P36981).

### Microscopy and image acquisition

Fluorescent images were acquired using a Nikon C2 laser scanning confocal microscope equipped with 405, 488, 561, and 647 nm lasers and a 60× Plan Apo oil-immersion objective lens (NA 1.4). Image acquisition was performed using NIS-Elements software in combination with a motorized stage. Images were captured in a 1024 × 1024 pixel format using tile scan mode with no overlap between tiles. To accommodate oil displacement between tile transitions, a brief pause of 500 msec was introduced after each stage movement. The Perfect Focus System (PFS) was employed throughout the acquisition to maintain consistent focus. Multiple fields were stitched using NIS-Elements to cover a sufficiently large region, including the entire ACC. Z-stacks were not acquired. Low-magnification images of whole brain sections were acquired using a Nikon ECLIPSE Ts2 inverted microscope equipped with a 2× Plan Fluor objective lens and a SONY α7 digital camera. All imaging parameters, including laser power, detector gain, and scanning speed, were kept constant within each experimental set to ensure comparability.

### Masking and ROI selection

Non-specific staining of vascular endothelium and the pia mater was occasionally observed when using anti-gephyrin (mouse IgG) in combination with anti-mouse IgG CF 633. To correct this, particles larger than 100 pixels were detected following thresholding, and identified region of interests (ROIs) were filled in black. Subsequently, to minimize the influence on histogram matching, ROIs were manually selected to exclude non-ACC regions as much as possible.

### Histogram matching

To select representative reference images for histogram matching, intensity histograms were computed from two-channel TIF images, and the Euclidean distance between each histogram and the median histogram across all images was calculated (Figure 2—figure supplement 1A). The five images with the smallest distances were automatically selected as candidate reference images. From these, a WT–VPA image pair was chosen in which both images had small Euclidean distances and were located in the same direction relative to the median. These images were used as references for histogram matching. All analyses were performed using Python 3.10 and the skimage.exposure module.

### Image processing and quantification

For particle detection and quantification, 16-bit dual-channel TIF images after histogram matching were first processed using a Laplacian of Gaussian (LoG) filter with channel-specific Gaussian blurring (σ = 2.4 for gephyrin channel; σ = 1.5 for PSD-95 channel). Processed images were divided into non-overlapping blocks (256 × 256 pixels), and each block was independently binarized using a fixed threshold. A distance transform and marker-based watershed algorithm were then applied to separate adjacent particles. Particles were filtered based on area (2–46 pixels) and circularity (0–1.0), and the following metrics were computed per block: total number of particles, total particle area, and mean particle area per particle:

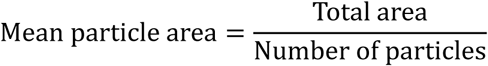

E/I ratio was calculated from raw values of excitatory and inhibitory synaptic density per mouse using the formula:

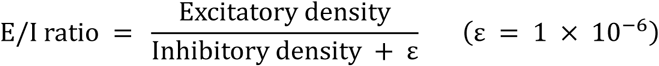

Instead of averaging individual E/I ratios, the mean E/I balance was derived by dividing the mean excitatory value by the mean inhibitory value.

To correct for optical distortion along the Y-axis (depth axis), a logarithmic inverse transformation was applied to each image after a user-defined 4-point perspective transformation. The corrected Y-coordinate *y*_orig_ corresponding to each rescaled coordinate *y*_new_ was computed as:

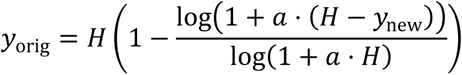

where *H* is the original image height and *α* is a distortion coefficient (typically 0.015). All image preprocessing, transformation, and batch quantification were performed using custom Python scripts with multiprocessing.

### Orientation analysis using structure tensor

To quantify local orientation, grayscale 8-bit TIF images were analyzed using a custom Python script based on structure tensor analysis, a method that estimates the predominant direction of local gradients by integrating spatial derivatives within a Gaussian-weighted neighborhood.

The input images were generated either by cropping ROIs from raw microscopy data or by applying geometric and logarithmic transformations to preprocessed images.

Image gradients were calculated using Gaussian derivative filters (σ = 3.0), and orientation angles (0– 180°) were computed from the structure tensor components as:

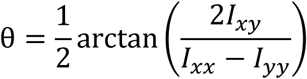

Anisotropy values, derived from the tensor components, were used as weights to generate orientation histograms. The global dominant orientation and circular standard deviation (SD) were estimated using vector summation with 2θ correction, and the results were visualized in a polar histogram with overlaid mean ± SD.

### Line regularity analysis

To assess the spatial and directional consistency of manually drawn lines, images were first geometrically corrected via perspective transformation and nonlinear y-axis compensation using the inverse log function:

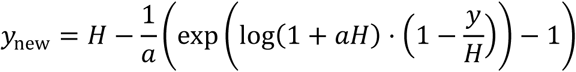

Coordinates from five reference lines across ten trials were extracted and fitted with linear regressions. Line spacing (intercepts) and direction (slopes) were analyzed using one-way ANOVA and Levene’s test to test for across-trial consistency.

### Laser microdissection

Mice were euthanized by cervical dislocation, and brains were rapidly removed and snap-frozen in isopentane cooled with liquid nitrogen. Frozen brains were embedded in optimal cutting temperature (OCT) compound (Sakura Finetek, Japan). Coronal sections (20 µm thick) were prepared using a cryostat (Leica CM3050 S) and mounted onto RNase-free polyethylene naphthalate (PEN) membrane slides (Leica Microsystems). Sections were fixed in cold methanol (−20 °C) for 3 minutes and then air-dried for 30 seconds using a cold air stream from a hair dryer. Laser microdissection of the ACC was performed using a Leica LMD7 system under bright-field guidance. Dissected tissue fragments were directly collected into the caps of RNase-free microtubes pre-filled with lysis buffer containing guanidine thiocyanate and reducing agents (from NucleoSpin® RNA Kit, Macherey-Nagel, Germany).

### Quantitative PCR

Total RNA was extracted from laser-microdissected ACC tissue using the NucleoSpin® RNA Kit according to the manufacturer’s instructions. RNA quantity and integrity were assessed using a TapeStation 4200 system (Agilent Technologies). Electrophoretic analysis showed no evident extra bands, although 18S and 28S peaks were approximately equivalent in size, indicating partial degradation (Figure 3—figure supplement 1D). Due to the extremely small amount of RNA recovered, RINe values could not be reliably obtained. All samples were subsequently used for qPCR analyses under these conditions. First-strand cDNA was synthesized using the PrimeScript™ FAST RT Reagent Kit with gDNA Eraser (Takara Bio, Japan), following the manufacturer’s protocol. Quantitative PCR was performed using a SYBR Green detection system on a StepOnePlus Real-Time PCR System (Applied Biosystems). Actb (β-actin) was used as an internal control. Gene-specific primer pairs used in this study are listed in Figure 3—figure supplement 1A. The relative position of each primer pair was calculated as the percentage distance from the 5’ end of the reference mRNA (RefSeq: NM_007393, NM_198862 and NM_007864), based on the midpoint of the amplicon. All reactions were run in triplicate, and gene expression levels were calculated using the ΔΔCt method on StepOne Software v2.3.

### Data analysis and statistical procedures

All statistical analyses and data visualizations were conducted in Python 3.10 using the following packages:

Data handling and computation: pandas 2.1.4, numpy1.26.3, scipy1.11.4

Statistical modeling: statsmodels 0.14.2, pingouin 0.5.5

Machine learning: scikit-learn 1.3.2

Plotting: matplotlib 3.8.2, seaborn 0.13.1

Quantification of histological parameters was conducted on three sections per mouse whenever available. Each mouse was represented by the average of these sections, yielding a single data point per animal (WT: *n* = 11; VPA: *n* = 11; WT Exercise: *n* = 3; VPA Exercise: *n* = 3). In some instances, due to tissue loss or suboptimal staining, only one or two sections per mouse were usable; these were included to maximize statistical power and were processed identically to complete datasets. As the unit of analysis was the individual mouse, and intra-animal variability was modest, partial inclusion was unlikely to bias group-level inferences.

### Group-level comparisons for behavioral and molecular measures

For comparisons of group-level differences in behavioral and molecular measures, Welch’s *t*-test was used due to its robustness to heteroscedasticity. However, pairwise *t*-test was used exclusively for the depth-wise analysis of layer 5 based on CTIP2 labeling (Figure 2E). Each parameter was tested independently without multiple-comparison correction across outcomes, as each assay represented a pre-specified, hypothesis-driven endpoint. Statistical significance thresholds were defined conventionally (*\*p* < 0.05, \*\**p* < 0.01, \*\*\**p* < 0.001), and selected *p*-values were annotated directly on the corresponding plots alongside the direction of group effects.

### Depth-wise correlation analysis and multiple testing correction

To examine correlations between behavior and synaptic architecture, we computed Pearson’s correlation coefficients between six behavioral measures and synaptic densities of PSD-95 and gephyrin. Synaptic values were averaged within five cortical depth bins (0–10%, 10–20%, 20–40%, 40–70%, 70– 100%) for each subject (*n* = 22). For hypothesis-driven analysis, we focused on the 10–20% bin for both markers, based on laminar enrichment (Figure 3C and D), resulting in 12 comparisons (6 behaviors × 2 markers). BH FDR correction (*α* = 0.05) was applied (Appendix 1—table 1). Exploratory correlations across all 60 combinations (6 behaviors × 2 markers × 5 bins) were also tested. Power analysis (two-tailed, *α* = 0.05, 1 − *β* = 0.8) indicated that only large effects (*|r|* ⪎ 0.55) could be reliably detected. Although several comparisons exceeded this threshold and reached nominal significance (*p* < 0.05), none remained significant after FDR correction (Appendix 1—table 2). These results are considered hypothesis-generating.

### Principal Component Analysis (PCA)

PCA was conducted on standardized (*z*-scored) behavioral and synaptic features to identify latent phenotypic dimensions across 22 mice (*n* = 11 WT, *n* = 11 VPA). Eight input variables were selected to represent excitatory and inhibitory synaptic densities, working memory (Y-maze), social interaction, locomotion, and anxiety-related behaviors. The first two principal components (PC1 and PC2) together explained ∼67% of the total variance. Based on feature loadings and vector orientations, PC1 was interpreted as reflecting excitatory synaptic structure (e.g., PSD-95), and PC2 as indexing anxiety-related behavior (e.g., center time in the open field test).

To evaluate the robustness of PCA decomposition, bootstrap resampling (*n* = 1,000) was performed. Axis alignment was enforced using two anchor variables PSD-95 density at 10% depth and open field center time to preserve the interpretability of PC1 and PC2. Axes were flipped when necessary to ensure consistent sign orientation. Loadings, explained variance ratios, and PC1–PC2 distributions were summarized to assess the stability of the component structure.

To assess whether PCA-transformed data space could reliably classify genotype groups, two classifiers—logistic regression and support vector machine (SVM) with an RBF kernel—were trained using the first eight PCA scores as input features. Five-fold stratified cross-validation was used to evaluate classification accuracy and area under the receiver operating characteristic curve (ROC AUC). To test the statistical significance of group separation, permutation testing was conducted with 1,000 label shuffles. For both classifiers, observed classification accuracy and permutation-derived *p*-values were reported.

To determine the optimal number of principal components, five-fold cross-validation was performed across 1 to 8 components (i.e., the total number of input variables). For each fold, reconstruction error was computed between the original and PCA-reconstructed test data. Mean squared error (MSE) was used as the performance metric, and results were visualized with error bars representing standard deviation.

### Linear discriminant analysis (LDA)

LDA was used to assess multivariate separation between WT and VPA-exposed mice based on behavioral and synaptic features. Input variables were standardized (mean = 0, SD = 1), and classification accuracy was estimated using leave-one-out cross-validation (LOOCV). For interpretability, LDA was also fitted to the full dataset to extract feature coefficients, indicating each variable’s contribution to group separation. LDA scores were visualized per subject with group-based color coding. To compare with univariate approaches, Welch’s *t*-tests were performed for each feature, and results were compiled alongside LDA coefficients.

### Multivariate profiling and clustering of VPA-resilient individuals

To assess group-level differences between VPA-resilient and non-resilient mice, Cliff’s delta (*Δ*) was calculated for eight predefined behavioral and structural features, including PSD-95 and gephyrin puncta density at 10% depth, social interaction time, and Y-maze performance. Confidence intervals were estimated via 1,000 bootstrap iterations. Only features with sufficient data (*n* ≥ 2 per group) were included. Results were visualized as error-bar plots displaying *Δ* with 95% confidence intervals.

To evaluate whether behavioral data alone could replicate multivariate resilience labels, we applied *k*-means clustering (*k* = 2) to two key behavioral features: social interaction time and Y-maze re-entry ratio. Clustering performance was quantified using the adjusted Rand index (ARI), comparing cluster assignments to *Z*-score-based labels. PCA was used to project the data into two dimensions for visualization. The ARI value was shown on the plot for reference.

Finally, cluster-level behavioral profiles were summarized by computing mean ± SEM for each variable. Grouped bar plots were used to compare clusters across the two behavioral features. These results supported visual interpretation of how behaviorally derived clusters relate to ASD-relevant phenotypes.

### Analysis of intervention effects

To evaluate the combined effects of genotype and exercise intervention, animals were grouped into four experimental conditions (WT Control, WT Exercise, VPA Control, VPA Exercise). For each behavioral or synaptic outcome, pairwise comparisons were conducted using the Games–Howell test, a variance-adjusted, nonparametric method robust to unequal sample sizes and variances. Effect sizes (Hedges’ *g*) were computed for all comparisons. Selected *p*- and *g*-values were annotated directly on bar plots to aid interpretation. No multiple-comparison correction was applied across these outcomes, as each endpoint corresponded to a distinct, independently hypothesized assay.

In addition, to assess exercise effects on laminar synaptic organization within the VPA-exposed group, depth-wise comparisons between VPA and VPA Exercise mice were conducted using the Games– Howell test. To adjust for repeated testing across depth bins, *p*-values were corrected using the BH FDR procedure. Group means ± SD were plotted across depth with significance markers indicating *q*-based thresholds.

### Validation of PCA stability after intervention inclusion

To ensure that the inclusion of exercise groups did not alter the underlying variance structure, we performed Procrustes analysis and cosine similarity comparisons between PCA configurations computed with and without the intervention data. These analyses confirmed that the overall geometry and principal component orientations of the PCA space were preserved.

## Funding

This work was supported by JSPS KAKENHI Grant Number JP24K09994 (to S.I.), JP22K06805 (to H.I.) and Chugai Foundation for Innovative Drug Discovery Science: C-FINDs (to S.I.).

## Author contributions: CRediT

M.M.: Conceptualization, Investigation (all experiments except for LMD and qPCR), Methodology, Formal analysis, Visualization, Writing – Original Draft

S.I.: Supervision, Funding acquisition, Formal analysis (PCA and LDA), Methodology, Software, Conceptualization, Investigation (LMD and qPCR), Validation, Visualization, Data curation, Writing – Original Draft, Writing – Review & Editing

H.I.: Project administration, Funding acquisition, Writing – Review & Editing, Validation, Resources

## Declaration of generative AI and AI-assisted technologies

We used ChatGPT for partial writing and revision of Python scripts, as well as for improving the readability and language of the manuscript during its preparation. After using ChatGPT, we reviewed and edited the content as necessary and take full responsibility for the final content of the published article.

## Declaration of Competing Interest

The authors declare that they have no known competing financial interests or personal relationships that could have appeared to influence the work reported in this paper.

## Acknowledgements

We thank all our lab members for technical advice. We thank Yoshihiro Morimura and Kyosuke Fukuda for technical assistance and care of the mice. We thank Kayo Suzuki (Education and Research Support Center, Gunma University) for the technical support for the qPCR and microscopic analysis. This work used equipment shared in the MEXT Project for promoting public utilization of advanced research infrastructure (JPMXS0420600124 and JPMXS0420600125).

## Data availability

The data supporting the findings of this study, including processed quantification tables and analysis scripts, are available on GitHub: https://github.com/sotaro-ichinose/Integrated-Profiling-of-Synaptic-Organization-and-Behavioral-Phenotypes

The dataset includes figure-linked ‘.csv’ files and analysis scripts in Python and ImageJ macro language. Raw imaging data are not publicly available due to size and ongoing analyses, but can be provided by the corresponding author upon reasonable request. Note that the dataset is associated with a manuscript currently under peer review, and may be revised prior to final publication.

---

**Figure 1—figure supplement 1.**
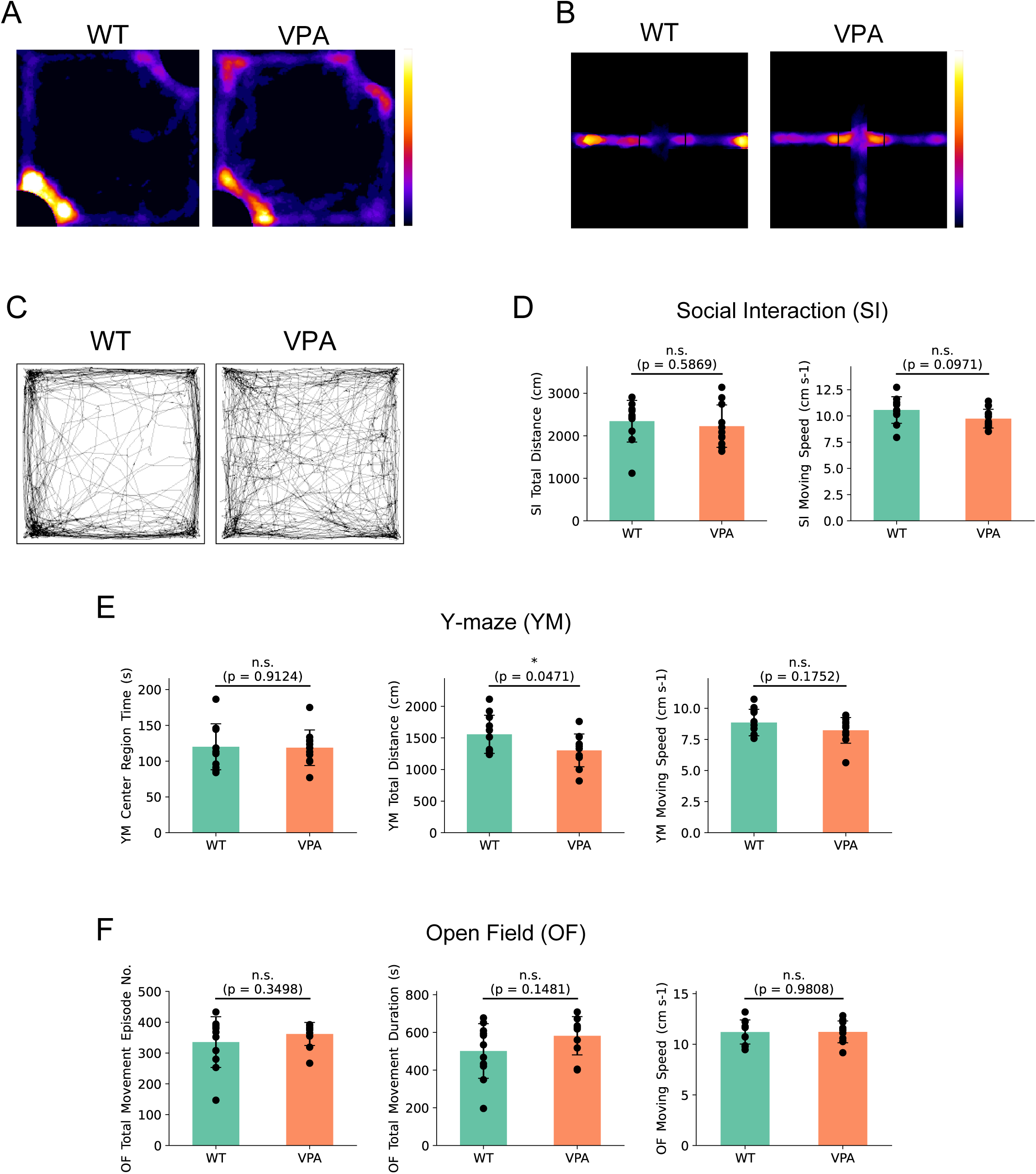
Behavioral analyses in WT and VPA-exposed mice. (A) Heatmaps of time spent during the SI test in WT and VPA mice (LUT: Fire; warmer colors indicate longer stay). (B) Heatmaps of time spent during the EPM test in WT and VPA mice (LUT: Fire). (C) Representative locomotor trajectories during the OF test. (D) Summary of SI test. Total moving distance and average moving speed. (E) Summary of YM test. Total time staying at center region, total moving distance and average moving speed. (F) Summary of OF test. Total movement episode number, totatl movement duration and average moving speed. Data are presented as mean ± SD. WT (*n* = 11), VPA (*n* = 11). Statistical analysis was performed using Welch’s *t*-test. * *p* < 0.05; ** *p* < 0.01; n.s. not significant.

**Figure 2—figure supplement 1.**
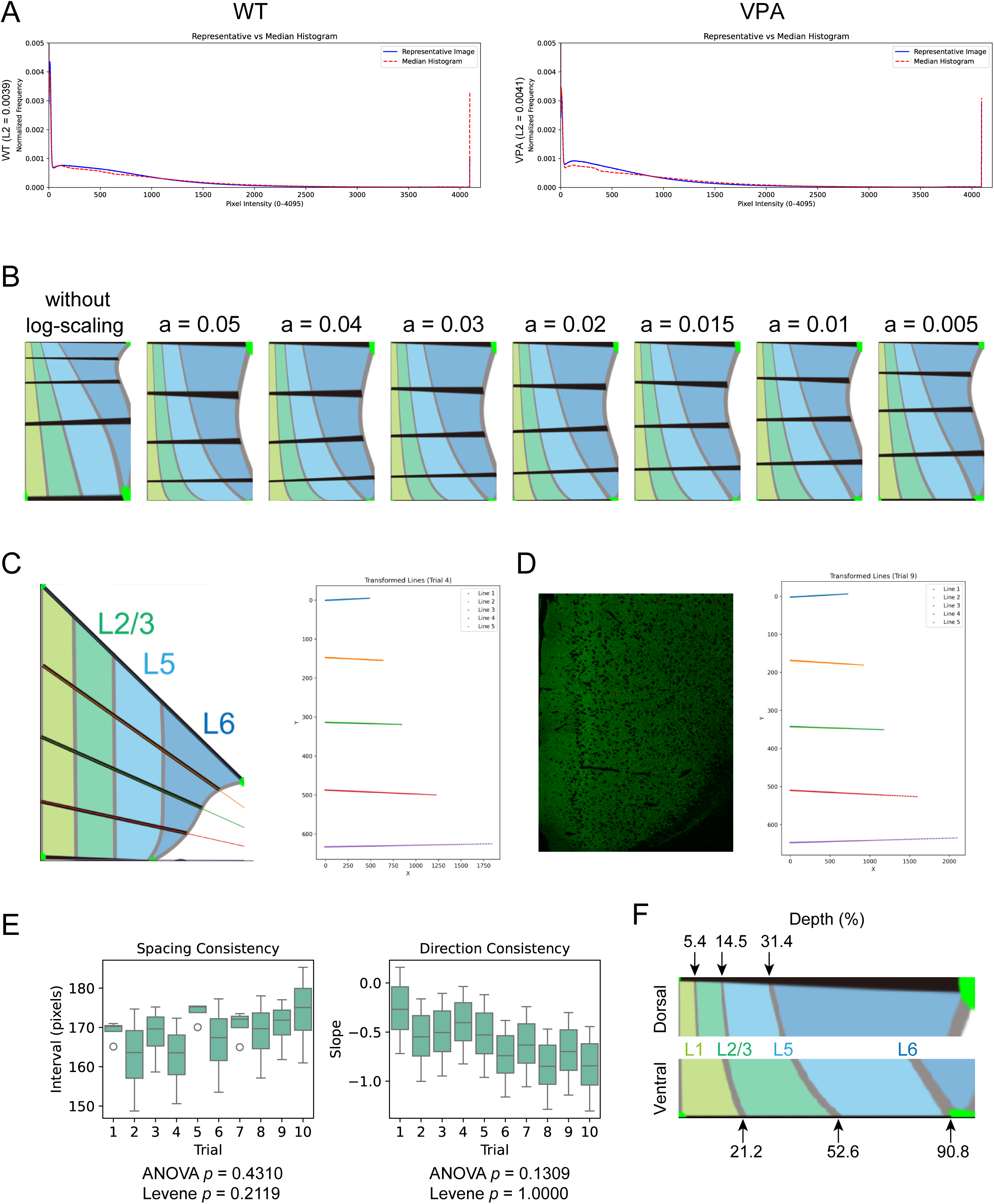
Perspective and log transformation for accurate depth-resolved analysis in the ACC. (A) Histograms of representative image and median pixel intensity from confocal images of the anterior cingulate cortex (ACC) in WT and VPA-exposed mice. The L2 distance between normalized intensity profiles in layer 2 was 0.0039 for WT and 0.0041 for VPA. Corresponding mean intensity differences were −71.6 and −93.8, respectively. All images were acquired in 12-bit format, with a maximum pixel value of 4095 representing the saturation level. (B) Effect of the constant parameter in logarithmic scaling on output image distribution. Without log-scaling, signal intensity is biased toward the ventral side, resulting in increased information density in that region. When the scaling constant was set to *a* = 0.015, information content became approximately uniform along the dorsal–ventral axis. (C) Model-based assessment of spatial uniformity and directional consistency after transformation. (D) Assessment of spatial uniformity and directional consistency after transformation using actual images. Five straight lines, defined as linear functions matching those in (C), were blindly placed within the image. Their transformed positions were then visualized to evaluate the effects of the transformation. (E) Spacing consistency and directional consistency were assessed across 10 repeated transformations. The test shown in panel D was performed 10 times, and statistical analyses were conducted. One-way ANOVA and Levene’s test revealed no significant differences (*p* > 0.05), indicating that both equidistance and directional uniformity were preserved across repeated applications of the transformation. *n* = 10 trials. (F) Difference in the transformed depth of cortical layers along the dorsal– ventral axis, assessed using a model diagram.

**Figure 3— figure supplement 1.**
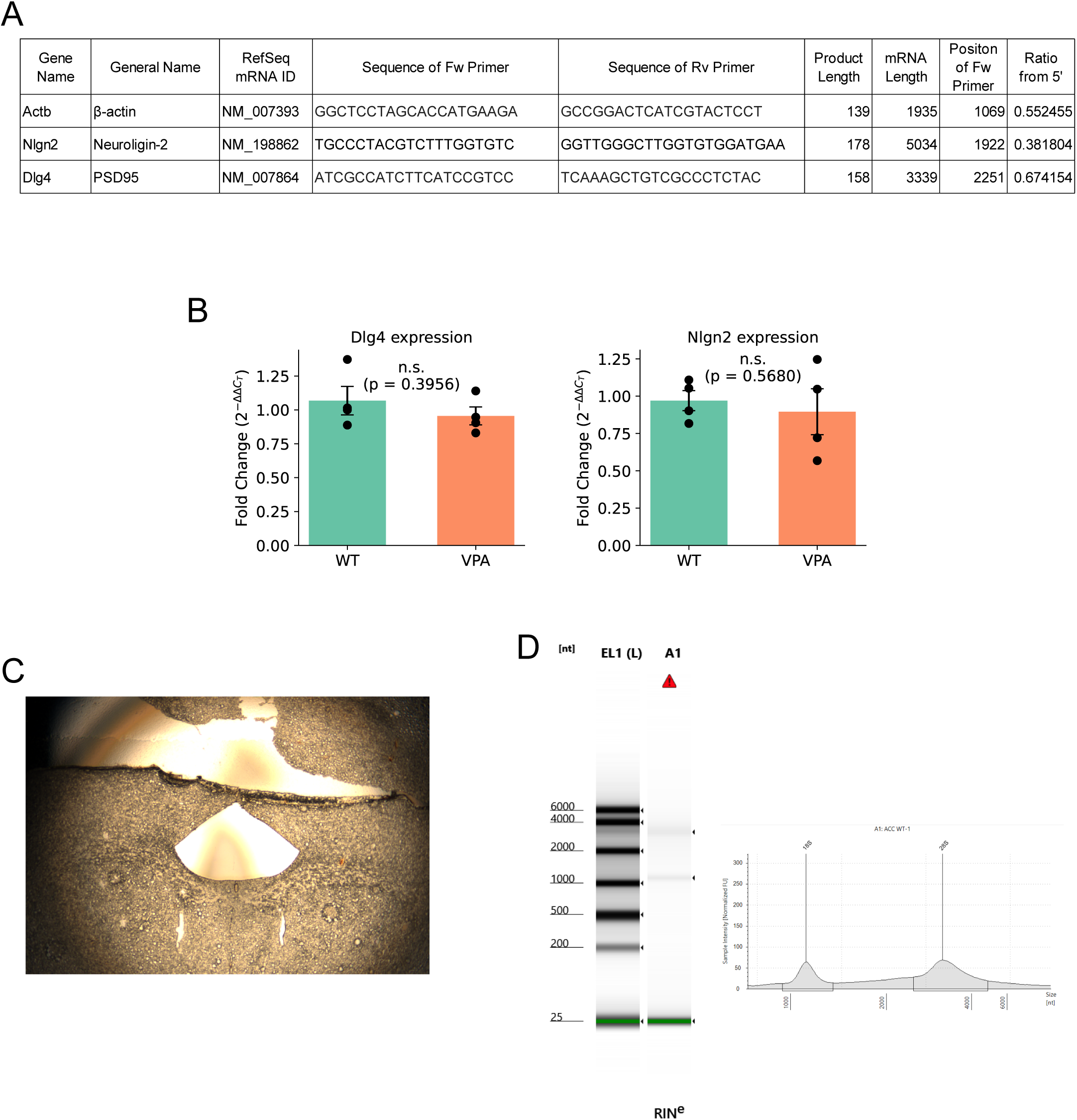
Quantification of synapse-related transcripts using LMD and qPCR. (A) Target genes and primer sequences used for qPCR. (B) Quantification results for Dlg4 and Nlgn2 analyzed by the ΔΔCt method, using Actb as the housekeeping gene. mean ± SD. Welch’s *t*-test. n.s., not significant. *n* = 4 mice, each with triplicate technical replicates. (C) Cortical image after LMD. (D) RNA quality assessment using TapeStation after RNA extraction. Due to the low amount of RNA, the RNA integrity number equivalent (RINe) could not be calculated.

**Figure 4— figure supplement 1.**
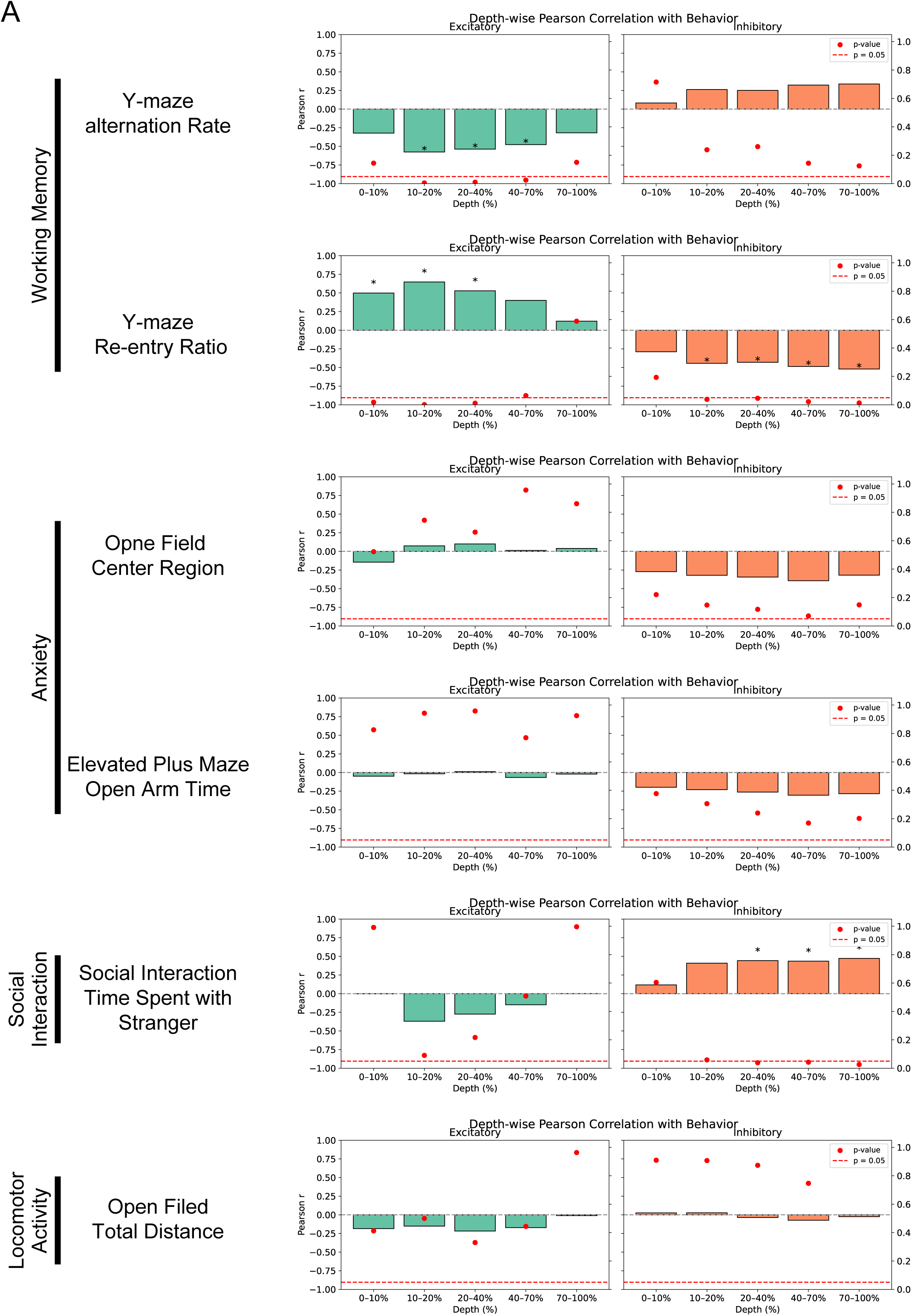
Uncorrected depth-wise Pearson correlations between behavioral performance and synaptic marker densities. Each bar represents the correlation coefficient (*r*) between behavioral scores and synaptic density averaged within each depth bin. Red dots indicate the corresponding *p*-values, and asterisks mark *p*-values prior to correction. These results are presented as exploratory analyses and should be interpreted statistically based on FDR-corrected values (Appendix1—table 1 and 2). *n* =22. * *p* < 0.05.

**Figure 4— figure supplement 2.**
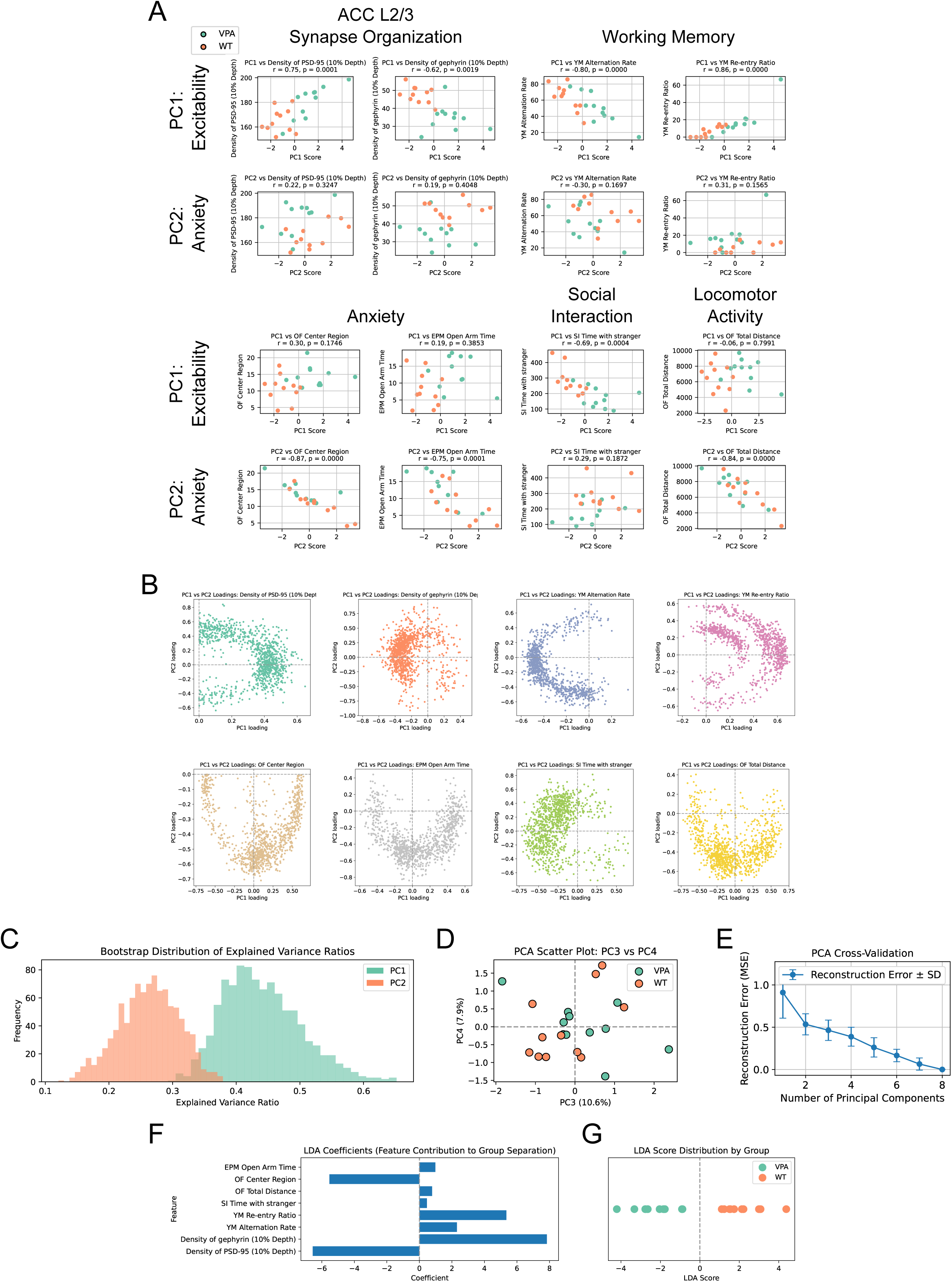
Multivariate integration of synaptic organization and behavioral analyses. (A) Correlations between each variable and principal components PC1 and PC2. Synaptic organization, working memory, and social interaction indices were strongly associated with PC1, while anxiety- and locomotion-related measures correlated more strongly with PC2. (B) Bootstrap distributions of variable contributions (*n* = 1000). All variables showed tightly clustered distributions, indicating high stability. (C) Bootstrap distributions of PC1 and PC2 variance explained (*n* = 1000). Both distributions were centered around PC1 = 36.6% and PC2 = 29.9%, demonstrating stable component structure. (D) Classification based on PC3 and PC4. These components contributed minimally to group separation, supporting the interpretability of PC1 and PC2. (E) Cross-validation analysis. No clear inflection point (“elbow”) was observed, indicating model robustness. (F) Linear discriminant analysis (LDA) coefficients. Both PSD-95 and gephyrin showed high coefficients, while the coefficient for social interaction—a core ASD-related phenotype—was relatively low. (G) LDA scores by group. WT and VPA mice were accurately classified based on the integrated dataset. For the all analyses, WT (*n* = 11), VPA (*n* = 11).

**Figure 5— figure supplement 1.**
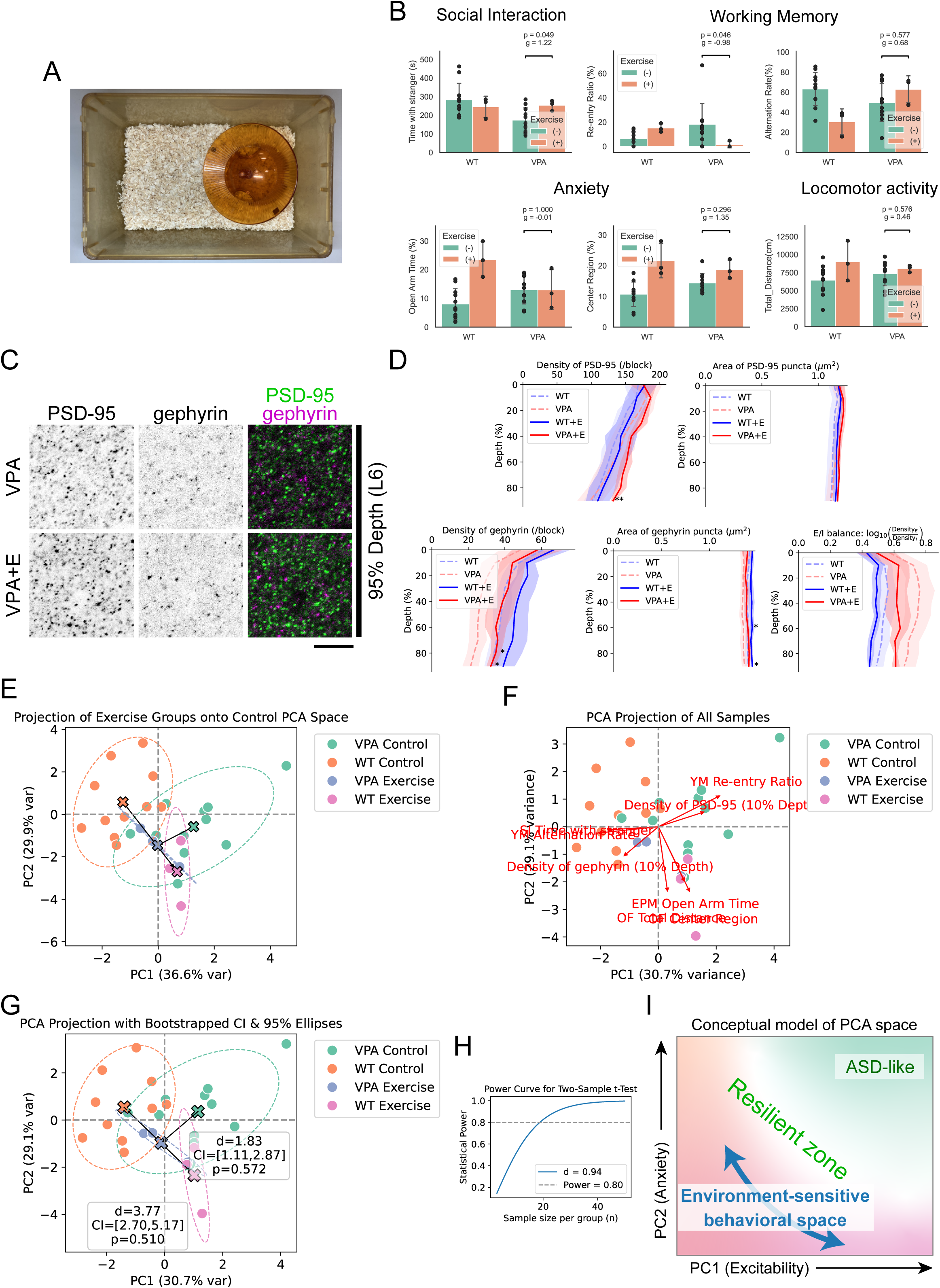
Application of the standardized pipeline to evaluate voluntary exercise intervention. (A) Photograph of the voluntary exercise apparatus. A mouse igloo and a running disk were placed inside the home cage. (B) Summary of behavioral outcomes. Comparisons are shown only for the VPA Control and VPA Exercise groups. mean ± SD. WT Control: *n* = 11; VPA Control: *n* = 11; WT Exercise: *n* = 3; VPA Exercise: *n* = 3. *p*-value (Games–Howell test) and effect size (Hedges’ *g*) for the VPA Control vs VPA Exercise comparison are shown directly on the bar graph for clarity. (C) Enlarged views of representative ACC regions at 95% depth, corresponding to cortical layer 6. Scale bar: 10 μm. (D) Summary of synaptic organization and E/I balance. Both PSD-95 and gephyrin densities showed upward trends following exercise, with a significant increase observed at 90% depth. Consequently, the E/I balance in the VPA Exercise group shifted closer to that of WT. Each block represents a 53.4 µm × 53.4 µm region. mean ± SD. WT Control: *n* = 11; VPA Control: *n* = 11; WT Exercise: *n* = 3; VPA Exercise: *n* = 3. *q* < 0.05, *q* < 0.01 by Games–Howell test with BH FDR correction. (E) Projection of WT Exercise and VPA Exercise data into the PCA space defined by Control groups. Centroids are indicated by × marks, and arrows denote their displacement. 95% CI ellipses are shown as dashed lines. (F) PCA recalculated using all animals. (G) Centroid shift analysis based on 1000 bootstrap resamples. The centroid of WT significantly shifted (Hotelling’s *T*² = 38.6, *p* = 0.0004), but non-parametric tests did not detect significant changes in WT or VPA following exercise. Centroids are indicated by × marks, and arrows denote their displacement. 95% CI ellipses are shown as dashed lines. (H) Power analysis curve. For an effect size of *d* = 0.94, sample sizes greater than *n* = 19 are required to achieve 80% power. (I) Conceptual model of PCA space. Integrating findings from resilient individual classification and exercise-induced shifts suggests that environmental sensitivity may extend across a broader region of PCA space. Under this view, ASD-like phenotypes may be more precisely confined to the upper-right quadrant, where both PC1 (excitability) and PC2 (anxiety) are elevated. Future studies with larger cohorts will be necessary to validate this model. For (E), (F) and (G), WT Control: *n* = 11; VPA Control: *n* = 11; WT Exercise: *n* = 3; VPA Exercise: *n* = 3.

## Table Legends

**Appendix 1—table 1 Behavior–synapse correlations with BH FDR correction (α = 0.05) applied across 12 comparisons (6 behaviors × 2 markers)** This table summarizes Pearson correlation coefficients between behavioral measures and synaptic marker densities (PSD-95 and gephyrin) across individual animals. Benjamini–Hochberg false discovery rate (BH FDR) correction was applied to adjust for multiple comparisons (*n* = 12). Statistically significant correlations are indicated in bold.

**Appendix 1—table 2 Behavior–synapse correlations with BH FDR correction (α = 0.05) applied across 60 comparisons (6 behaviors × 2 markers × 5 bins)** This table summarizes Pearson correlation coefficients between behavioral measures and synaptic marker densities (PSD-95 and gephyrin) across individual animals. BH FDR correction was applied to adjust for multiple comparisons (*n* = 60). Statistically significant correlations are indicated in bold.

**Appendix 2—table 1 Classification performance of logistic regression and SVM (RBF kernel) including permutation test results** Mean accuracy and ROC AUC values were computed via cross-validation. Permutation tests (*n* = 1000) confirmed that observed accuracies were significantly higher than chance (permutation *p* < 0.001).

**Appendix 3—table 1 Group-wise comparison and linear discriminant analysis (LDA) of synaptic and behavioral features** This table summarizes group-level comparisons between WT and VPA-exposed mice across synaptic (gephyrin and PSD-95 densities at 10% depth) and behavioral features. Welch’s *t*-tests were used to evaluate group differences (*p*-values shown), with directionality indicated in the “WT > VPA” column. LDA coefficients reflect each variable’s contribution to group classification. The “LDA suggests WT > VPA” column indicates whether the direction of the LDA-based separation was consistent with the group mean comparison.

**Appendix 4—table 1 Hotelling’s *T*² test comparing WT Control and WT Exercise groups** This table shows the result of a Hotelling’s *T*² test assessing multivariate differences between the WT Control and WT Exercise groups across two principal components (PC1 and PC2). The test yielded a significant group difference (*T*² = 37.95, *F*(2, 11) = 17.39, *p* = 0.00039), indicating that voluntary exercise induced a significant shift in multivariate PCA space within the WT group.

**Appendix 4—table 2 Hotelling’s *T*² test comparing VPA Control and VPA Exercise groups** This table reports the result of a Hotelling’s *T*² test evaluating multivariate differences between the VPA Control and VPA Exercise groups based on PC1 and PC2 scores. No significant difference was observed (*T*² = 3.38, *F*(2, 11) = 1.55, *p* = 0.255), indicating that the exercise intervention did not produce a statistically detectable shift in multivariate PCA space for VPA-exposed mice under the current sample size.

## Notes

### Competing Interest Statement

The authors have declared no competing interest.

